# Barley HvNIP2;1 aquaporin permeates water, metalloids, saccharides, and ion pairs due to structural plasticity and diversification

**DOI:** 10.1101/2023.04.17.537278

**Authors:** Akshayaa Venkataraghavan, Hoshin Kim, Julian G. Schwerdt, Alexey V. Gulyuk, Abhishek Singh, Yaroslava G. Yingling, Stephen D. Tyerman, Maria Hrmova

## Abstract

Aquaporins can facilitate the passive movement of water and small polar molecules and some ions. The barley Nodulin 26-like Intrinsic Protein (HvNIP2;1) embedded in liposomes and examined through stopped-flow light scattering spectrophotometry and *Xenopus* oocyte swelling assays was found to permeate water, boric and germanic acids, sucrose and L-arabinose but not D-glucose or D-fructose. Other saccharides, such as neutral (D-mannose, D-galactose, D-xylose, D-mannoheptaose) and charged (N-acetyl D-glucosamine, D-glucosamine, D-glucuronic acid) aldoses, disaccharides (lactose, cellobiose, gentiobiose, trehalose), trisaccharide raffinose, and urea, glycerol, and acyclic polyols were permeated to a much lower extent. Apparent permeation of hydrated KCl and MgSO_4_ ion pairs was observed, while CH_3_COONa and NaNO_3_ permeated at significantly lower rates. Experiments with boric acid and sucrose revealed no apparent interaction between solutes when permeated together, and AgNO_3_ blocked the permeation of all solutes. Full-scale steered molecular dynamics simulations of HvNIP2;1 and spinach SoPIP2;1 revealed possible rectification for water, boric acid, and sucrose transport, and defined key residues interacting with permeants. In a biological context, the simulated sucrose rectification could mediate its apoplastic-to-intracellular transport but not the reverse, thus, constituting a novel element of plant saccharide-transporting machinery. Phylogenomic analyses of 164 Viridiplantae and 2,993 Archaean, bacterial, fungal, and Metazoan aquaporins rationalised solute poly-selectivity in NIP3 sub-clade entries and suggested that they diversified from other sub-clades to acquire a unique specificity of saccharide transporters. Solute specificity definition in NIP aquaporins could inspire developing plants for sustained food production.

**Significance Statement:** Aquaporins are fundamental to water and solute movements in nearly all living organisms. Solute selectivity inspections of the HvNIP2;1 aquaporin revealed that it transported water, hydroxylated metalloids boric and germanic acids, sucrose, L-arabinose, KCl, and MgSO_4_ ion pairs, but not D-glucose or D-fructose and to lesser extent urea, and acyclic polyols. This poly-selective transport by HvNIP2;1 classified in the NIP3 sub-clade aquaporins may afford nutritional and protective roles during plant development and in response to abiotic stresses. It is anticipated that the solute specificity definition of HvNIP2;1 inspires protein engineering and in silico mining to develop plants, which when exposed to suboptimal soil conditions of high soil metalloids, would overcome toxicity for sustained food production.

## Main Text

### Introduction

Aquaporins (AQPs) are fundamental to water and other solute movements in nearly all living organisms. AQPs are membrane proteins classified in the family of major intrinsic proteins (MIPs) that transport water and neutral solutes (1, 2), although the repertoire of permeated solutes has recently expanded as some AQPs may also transport ions (3). The AQP family consists of seven sub-families of plasma membrane intrinsic proteins (PIPs), tonoplast intrinsic proteins (TIPs), Nodulin 26-like Intrinsic Proteins (NIPs), and small and basic intrinsic proteins (SIPs). Other less common sub-families include Solanaceae X-intrinsic proteins (XIPs) (4) found in fungi, slime moulds and dicots, while GlpF-like intrinsic proteins (GIPs) (5), and hybrid (between PIPs and TIPs) intrinsic proteins (HIPs) are found in green algae, the moss *Physcomitrella* and lycopods (4). AQPs from each sub-family differ in physicochemical properties that underlie their complex roles in osmotic homeostasis, root water uptake from the soil, root and leaf hydraulic conductance, lateral root emergence, motor cell movement, internode elongation, the diurnal regulation of leaf movements, petal development, biotic interactions, signalling and stomatal dynamics (3, 6, 7).

NIPs, named after soybean nodulin 26 (NOD26), an abundant protein in the peribacteroid membrane of nitrogen-fixing root nodules, are functional equivalents of aquaglyceroporins (8), and occur in barley (9), rice (10), wheat (11) and other plants. Besides water and boric acid (BA) (9, 10) NIPs transport other hydroxylated metalloids such as arsenious and germanic acid (9,12–15), glycerol (12), urea (16), hydrogen peroxide (13), acyclic polyols, purines and pyrimidines (17), neutral lactic acid (18), selenious and antimonious acids (19) and aluminium malate (20). Some AQPs facilitate gas diffusions such as CO_2_ (21), NH_3_ (22–24) and O_2_ (25). When NOD26 was incorporated in lipid bilayers, ionic currents were detected with slight anion over cation selectivity (26, 27). From an evolutionary point of view, plant NIPs (28) and Solanaceae XIPs (‘metalloido-porins’) (29), permeating heavy metals and H_2_O_2_ but not water (30), were shown to evolve from cyanobacteria (31) some 1,500 million years ago. Then a primordial AQP NIP-like (*aqpN*) gene was acquired by some plants *via* horizontal transfer shedding light on NIPs solute evolution and selectivity (32, 33).

The structural monomeric sub-unit of functional AQP folds into six tilted membrane-spanning α-helices and two short re-entrant α-helices, running in two repeats, with five interconnecting loops forming a right-handed α-helical bundle. A permeating pore running through each monomer has a defined morphology and width in each AQP sub-family; for example, barley HvNIP2;1 has a wide pore along its entire length compared to PIPs and TIPs (2, 34, 35). In native environments, AQPs oligomerize into tetramers along a four-fold rotational axis and create a tightly fitted trapezoid that at structural and functional levels may form an additional central fifth pore. Each AQP monomer has its solute path and with a central pore, the tetramer potentially offers five permeating routes (3, 36). The broad solute selectivity of the monomer pore of NIPs is supported by Asn-Pro-Ala (NPA) motifs that are separated by 4-5 Å from the aromatic/R selectivity filter residues (2, 36).

To achieve the constant adjustment of water and solute permeability in fluctuating environments, cells developed multiple controls of AQP function through structural features. For example, extended N-and C-terminal regions in NIPs (9, 37) could impact transport, gating and *in vivo* AQP expression levels. These regions and cytosolic loops (*e.g.* loop D) in SoPIP2;1 (PDB 2B5F) (38, 39), AtPIP2;4 (PDB 6qim) (40), AtTIP2;1 (PDB 5i32) (41) and recently elucidated rice OsNIP2;1 (PDB 7nl4 and 7cjs) (42, 43) have similar spatial dispositions suggesting that dynamics of PIPs, TIPs, and NIPs could be similar. In SoPIP2;1 loop D alongside His193 and dephosphorylated Ser residues trigger pore closure and other structural re-arrangements (38).

To understand the function of HvNIP2;1 in the context of metalloid toxicity it is fundamental to understand that this trait is a major problem in cereal crops around the world (44). Although metalloids boron and silicon are essential micronutrients, other metalloids such as arsenic and germanium are toxic at certain concentrations, which together with high soil boron pose risks to populations (45). Metalloids exist in soils in the form of amphoteric oxides with atomic radii between 3.43 and 4.48 Å, and these differences could be exploited to engineer mutants with selective permeation properties (19).

In the present study, solute selectivity is explored in HvNIP2;1 solubilised from *Pichia pastoris* membranes through embedding in liposomes. The selectivity of HvNIP2;1 towards neutral solutes and ion pairs is examined, establishing that this AQP permeated water, BA, germanic acid, sucrose, and MgSO_4_ and KCl ion pairs at relatively high rates, but also permeated at low rates acyclic polyols, urea and glycerol, other ion pairs, and some mono-and disaccharides of various stereo chemistries. Steered molecular dynamics (steered MD) simulations were performed on the monomeric HvNIP2;1 3D model to provide atomic-level descriptions of permeation dynamics, including identifying structural elements that underlie solute poly-specificity. These data are compared to those of water-selective SoPIP2;1 AQP in an open state conformation (38, 39). Structural data suggested that the wider pore of HvNIP2;1 lined with acidic and aromatic residues, and high pore flexibility, were the key features supporting the permeation of a wide range of solutes. These data were rationalised by phylogenomic analyses of 3,157 Viridiplantae (green algae and land plants), Archaean, bacterial, fungal, and Metazoan AQPs to show that the members of the NIP3 sub-clade evolved a unique solute specificity of saccharide transporters through neo-functionalisation after the emergence of tracheophytes. The HvNIP2;1 substrate poly-specificity definition underscores yet unidentified roles of this AQP in plant growth, metabolism, and development.

## Results

### Cloning, expression, solubilisation and liposomal reconstitution of HvNIP2;1

The native *HvNIP2;1*-Myc-6xHis-pPICZ-B DNA fusion (46–48) was expressed in *P. pastoris*, solubilised in styrene-maleic anhydride co-polymer 3:1 (SMA) (49) and purified by Immobilised Metal Affinity Chromatography (IMAC) (46, 47). Near-homogenous HvNIP2;1 (Fig. 1A) was reconstituted in 1, 2-dimyristoyl-sn-glycero-3-phosphocholine (DMPC) liposomes that were floated on an iohexol (Accudenz) gradient to obtain homogenous populations of proteo-liposomes of around 100 μm in size (Fig. 1B) (47), which were used to define permeation properties of HvNIP2;1. Further details on cloning, expression, solubilisation, purification, and liposomal reconstitution of HvNIP2;1 are specified in Supplementary Information.

**Fig. 1.**
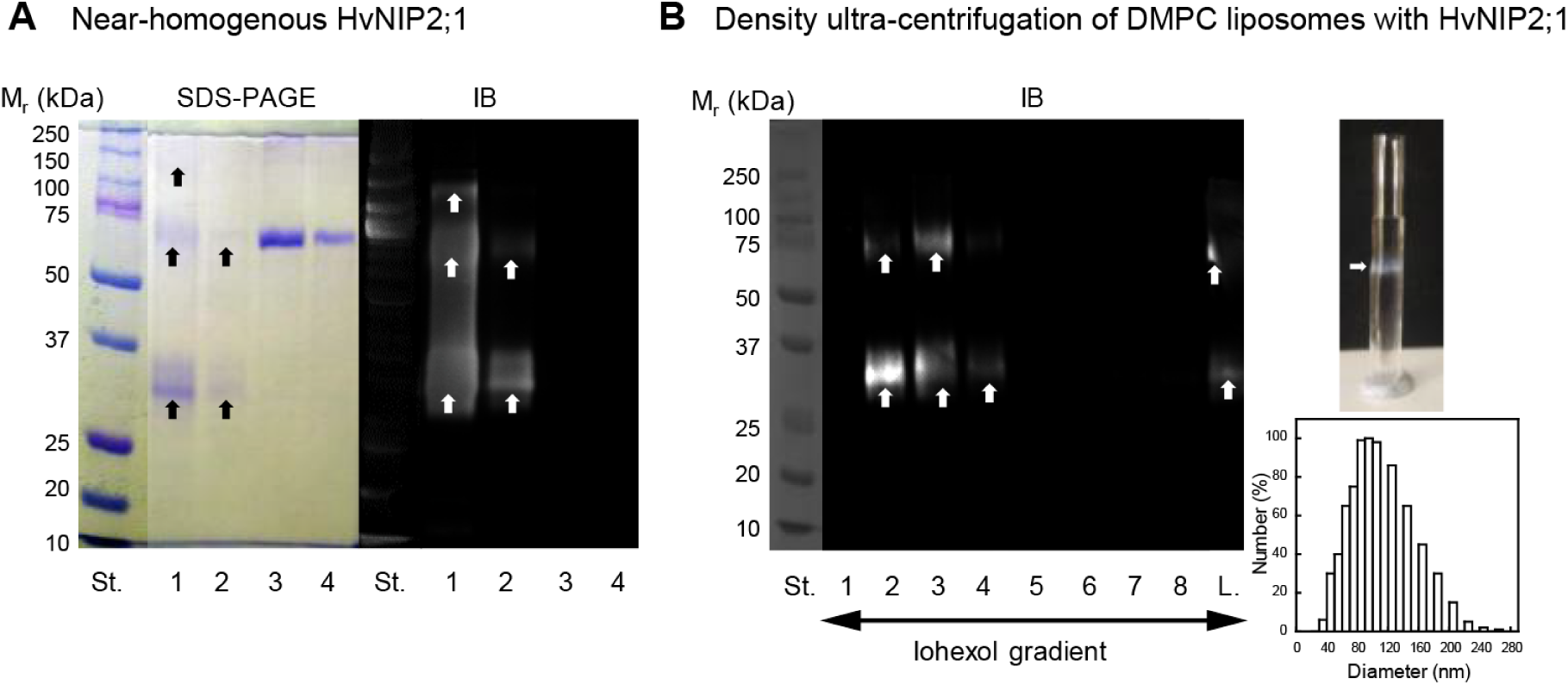
SDS-PAGE and immunoblot (IB) analyses of near-homogenous HvNIP2;1 purified by IMAC, and density gradient ultra-centrifugation of DMPC liposomes with reconstituted HvNIP2;1. *(A)* Monomeric (apparent molecular mass 34 kDa), dimeric (70 kD), and tetrameric (150 kDa) forms of HvNIP2;1 (lanes 1 and 2) are indicated by arrows at approximately 0.5 and 1 µg protein loadings, respectively. Lanes 3 and 4 contain BSA (Fraction V) at 1 or 2 µg loadings, respectively. Immunoblot (IB) analysis proceeded with anti-His antibody. St. lanes indicate molecular masses of protein standards. *(B)* Density gradient ultra-centrifugation of DMPC liposomes with embedded HvNIP2;1 and proteo-liposomal sizing. Lanes 1-8 are fractions collected after ultra-centrifugation in the iohexol gradient, where L. (lane 9) contains non-fractionated preparation. St. lane indicates molecular masses of protein standards. HvNIP2;1 was detected by IB with an anti-His antibody. Right top panel shows the test tube of liposomes with reconstituted HvNIP2;1 (arrow) after iohexol gradient floating, forming a white diffuse band. Bottom image displays the size distribution profile of proteo-liposomes analysed by NICOMP 380 Particle Sizing System.

### Permeation properties of HvNIP2;1

Measurements of water and solute permeability, based on osmotic gradients generated by co-incubating DMPC liposomes with embedded HvNIP2;1 or control liposomes lacking the protein, and osmolytes at high concentrations, were assessed by stopped-flow light scattering spectrophotometry (Fig. 2; Figs. S1 and S2; Table S1). These co-incubations created an osmotic gradient that in the first phase led to increased rates of water efflux causing the shrinkage of liposomes, while in the second phase, proteo-liposomes increased their volumes and swelled due to rate-limiting solute transport of water (50). This occurred with liposomes with reconstituted HvNIP2;1 that mediated solute permeation, but not for control liposomes or when liposomes with embedded HvNIP2;1 were incubated with 0.5 mM AgNO_3_, a potent blocker of AQPs (51), for which a re-swelling phase was incomplete (Fig. 1C). HvNIP2;1 permeated BA and germanic acid at higher rates, while glycerol, D-mannitol and D-sorbitol, and urea were permeated at low rates (Fig. 2; Figs. S1 and S2; Table S1). Surprisingly, HvNIP2;1 permeated at high rates the disaccharide sucrose, as previously observed (46), and monosaccharide L-arabinofuranose, and hydrated ion pairs KCl and MgSO_4_, while lactose, the CH_3_COONa and NaNO_3_ ion pairs were transported at lower rates. Conversely, NaF, the neutral monosaccharides D-xylopyranose, D-glucopyranose, D-fructopyranose, D-galactopyranose, D-mannopyranose and D-mannopyranopheptaose, charged monosaccharides D-glucosamine, N-acetyl β-D-glucosamine and D-glucuronic acid, disaccharides trehalose, cellobiose and gentiobiose, and trisaccharide raffinose were permeated at even lower rates (Fig. S2). Permeation rate constants and derived permeability coefficients (*P*) (Table S1) for water, metalloids, saccharides, and other solutes, and ion pairs descended in these orders:

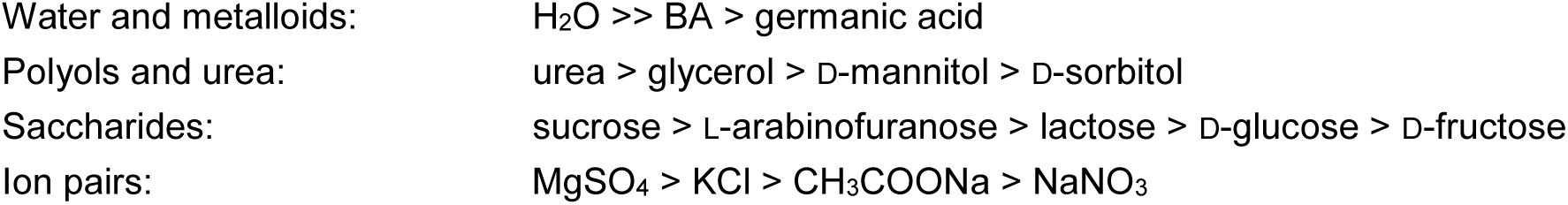
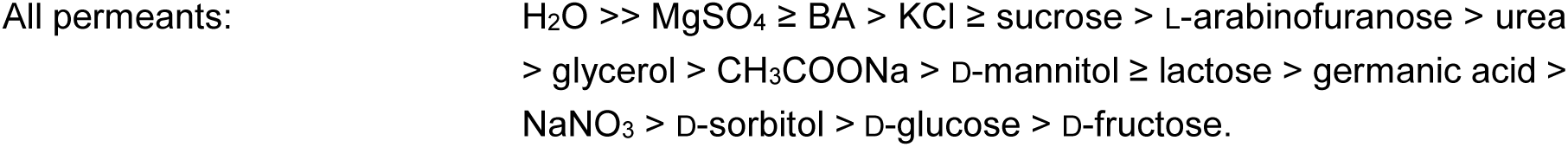

**Fig. 2.**
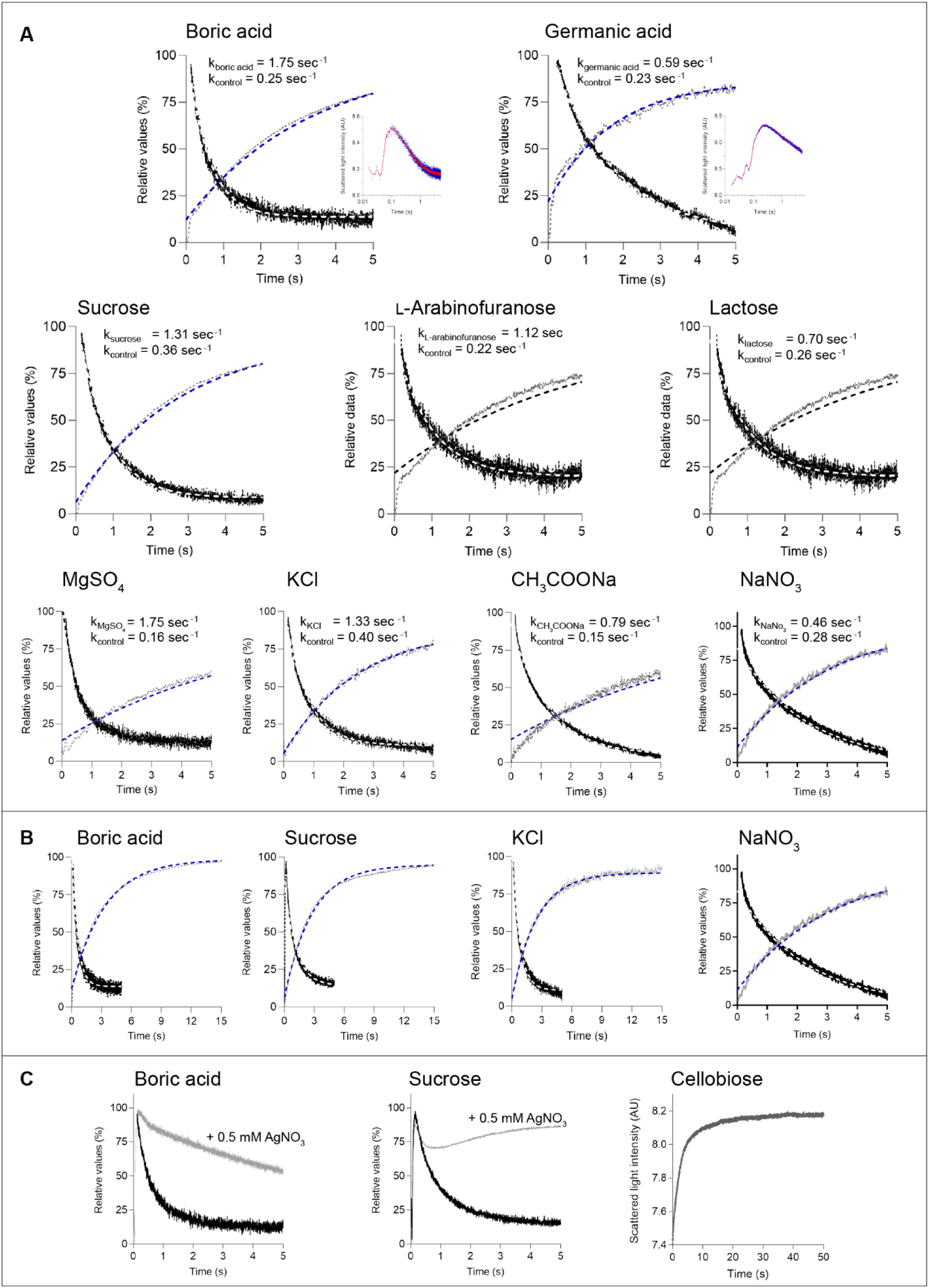
Transport of permeants by HvNIP2;1 embedded in liposomes. *(A-C)* HvNIP2;1 solubilised by SMA and reconstituted in DMPC liposomes were exposed to gradients of solutes generating osmotic gradients with BA and germanic acid (top panels), where insets show full profiles (shown also in Fig. S1), monosaccharide L-arabinofuranose and disaccharides sucrose and lactose (bottom-middle panels), and MgSO_4_, KCl, and NaNO_3_ (bottom panels). The uptake of permeants was measured by stopped-flow spectrophotometry. The light scattering traces indicate swelling slopes of HvNIP2;1 proteo-liposomes, where black curves indicate normalised data (in relative values) and white dashed curves represent non-linear fits using a one-phase decay model. Light scattering traces of control liposomes (lacking HvNIP2;1) are drawn in grey for the first five (panels A) and 15 sec (panels B), and non-linear fits using a one-phase association model are shown in blue dashed curves. 0.5 mM AgNO_3_ was used to inhibit transport in HvNIP2;1 (panels C; grey curves). In panel A, calculated rate constants for each combination of permeants are indicated in s^-1^ for liposomes with HvNIP2;1 and control liposomes lacking HvNIP2;1. Plots in panels B (control liposomes and proteo-liposomes for BA, sucrose, KCl, and NaNO_3_) and panel C (right plot – data of control liposomes exposed to cellobiose) indicate that no solute leakage was observed; the same non-solute leakage profiles were observed for control liposomes. Data were plotted in relative values (%) in GraphPad Prism 9.

Measurements of water and solute permeability for selected solutes by HvNIP2;1 were confirmed with *X. laevis* oocytes that were injected with cRNA of HvNIP2;1 or water as controls (Fig. 3), where the swelling was measured in response to the BA, sucrose, and D-glucose concentration gradients (9). Transport of BA and sucrose was observed, while D-glucose permeation did not occur (Fig. 3A) as shown using stopped-flow light scattering analyses (Fig. S2). No detectable interaction was seen between BA and sucrose when permeated together (Fig. 3B). In oocytes, incubated with 0.5 mM AgNO_3_ (50, 51) – the known histidine and thiol groups modifier, all solute transport was blocked.

**Fig. 3.**
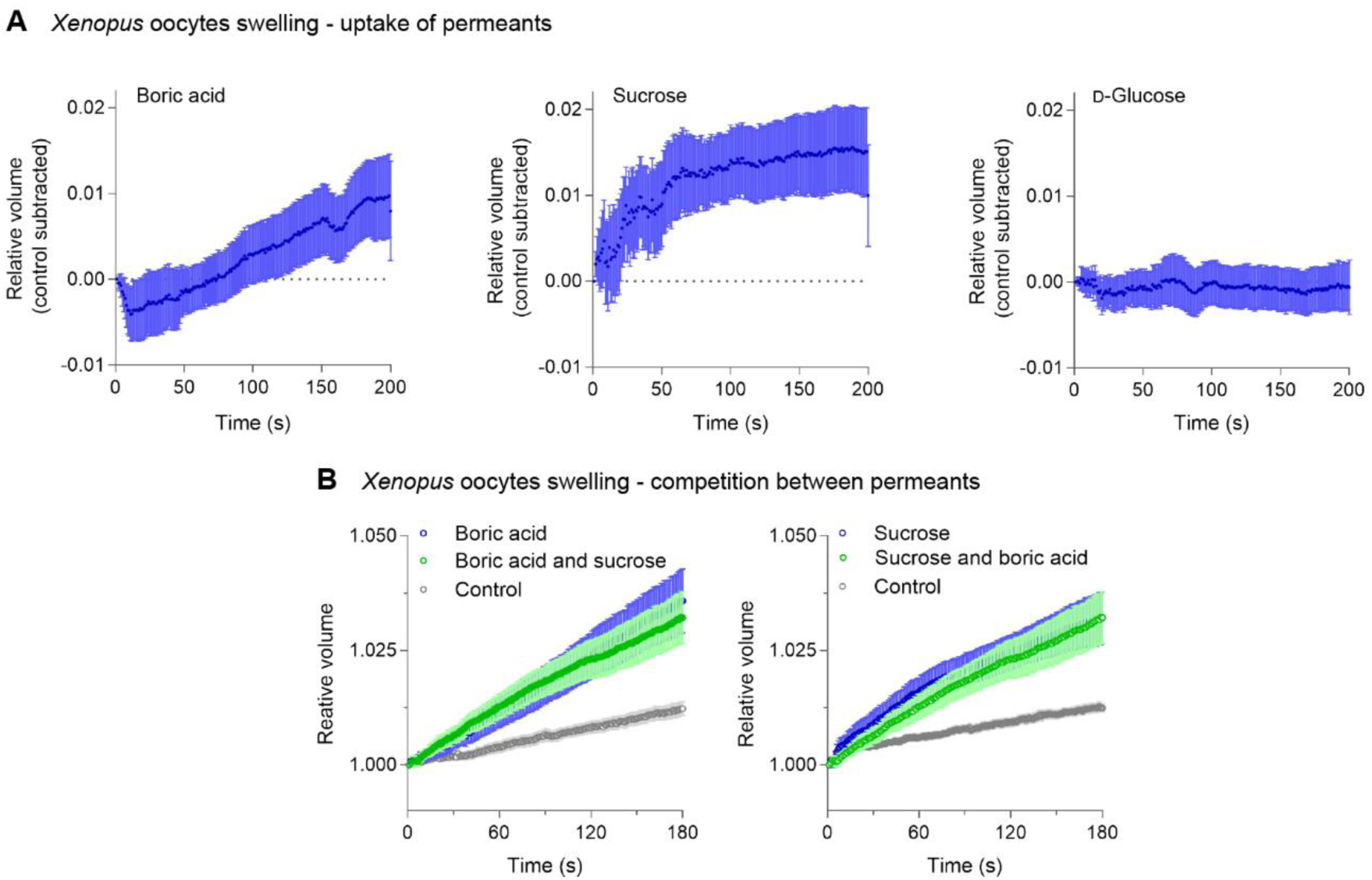
Uptake of permeants by HvNIP2;1 expressed in *Xenopus oocytes*. *(A)* Transport of 160 mM BA, sucrose, and D-glucose by oocytes transformed by cRNA. An increase in relative volume (V/V_0_; control subtracted) of oocytes after replacing incubation solutions with solutions supplemented with 160 mM BA, sucrose, and D-glucose is shown. *(B)* Transport 160 mM BA and sucrose alone or together (each in 80 mM concentrations) in *X. laevis* oocytes. Controls represent water-injected oocytes subjected to solutes, as described in panel A.

### Molecular model of HvNIP2;1 and the SoPIP2;1 crystal structure

The 3D model of HvNIP2;1 (with 94% confidence level at an accuracy higher than 90%) generated by homology modelling, as described in Supplementary Information, featured six tilted membrane-spanning α-helices, two short re-entrant α-helices in two repeats, and five interconnecting loops forming an α-helical bundle (Fig. S3A). As observed, individual monomers formed a quaternary assembly (Fig. 1A), and solute-conducting pore, running through each monomer in HvNIP2;1, were wide alongside their lengths without forming constricted regions observed in other AQPs (Fig. S3). Dispositions of α-helices in HvNIP2;1 followed a pseudo-two-fold axis that runs along a membrane normal and sub-divided an hour-glass fold into bipartite segments. A significant symmetrical distribution of repeating peptide motifs was observed in each bipartite HvNIP2;1 segment (Fig. S3B; Table S2) detected by the Multiple EM (Expectation Maximization) for Motif Elicitation (MEME) analysis (52). At least ten peptide motifs were identified in each segment based on P-values (Table S2).

The 3D model of HvNIP2;1 was relaxed using the all-atom MD simulation and used in steered MD simulations, where the force with a spring constant of 0.1 kcal/mol/Å_2_ and a pull rate of 0.05 fN per 2 fsec simulation time-step in a pore direction was applied to solutes along a chosen direction, to determine transport competencies and rates to understand water, metalloid, saccharide, and ion pair permeation mechanisms. To increase the accuracy of data interpretation, the monomeric high-efficiency water-permeating SoPIP2;1 AQP in an open state conformation (38, 39) was used as an accessory structural target in steered MD simulations.

### Steered MD simulations of solute permeation by HvNIP2;1 and SoPIP2;1

Initial steered MD simulations (53, 54) of solute permeation, and evaluations of pore volumes and structural features allowed us to form hypotheses. These and further MD simulations (Figs. 4-7; Figs. S4-S10) indicated that SoPIP2;1 transported water faster than HvNIP2;1 (average times for one molecule 3.7-4.7 nsec for SoPIP2;1 *versus* 5.1 to 8.3 nsec for HvNIP2;1) under the equivalent force, but with a lower total number of water molecules (maximum 7.2-9.3 for SoPIP2;1 *versus* 11.4-13.7 for HvNIP2;1) (Fig. 4A). A significantly more voluminous pore of HvNIP2;1 (5,100 Å_3_) compared to that of SoPIP2;1 (2,500 Å_3_) (Fig. S5) accommodated a higher number of water molecules, but the ‘X-shaped’ pore shape (Fig. 4; Fig. S5) generated a bottleneck that subjugated water permeability. Bearing in mind the importance of bi-directionality and pore shapes of AQPs during transport, permeation directionalities, designated A-B-C (A-entrances are at cytoplasmic sides of AQPs) and C-B-A (C-entrances are at extracellular or apoplastic sides of AQPs) were considered (Fig. S5). Here, A-B-C (pointing from cytoplasmic to apoplastic) and C-B-A (from apoplastic to cytoplasmic) directionalities or paths were defined as all-through permeation modes, while A-B-A and C-B-C were deemed to be U-turn modes.

**Fig. 4.**
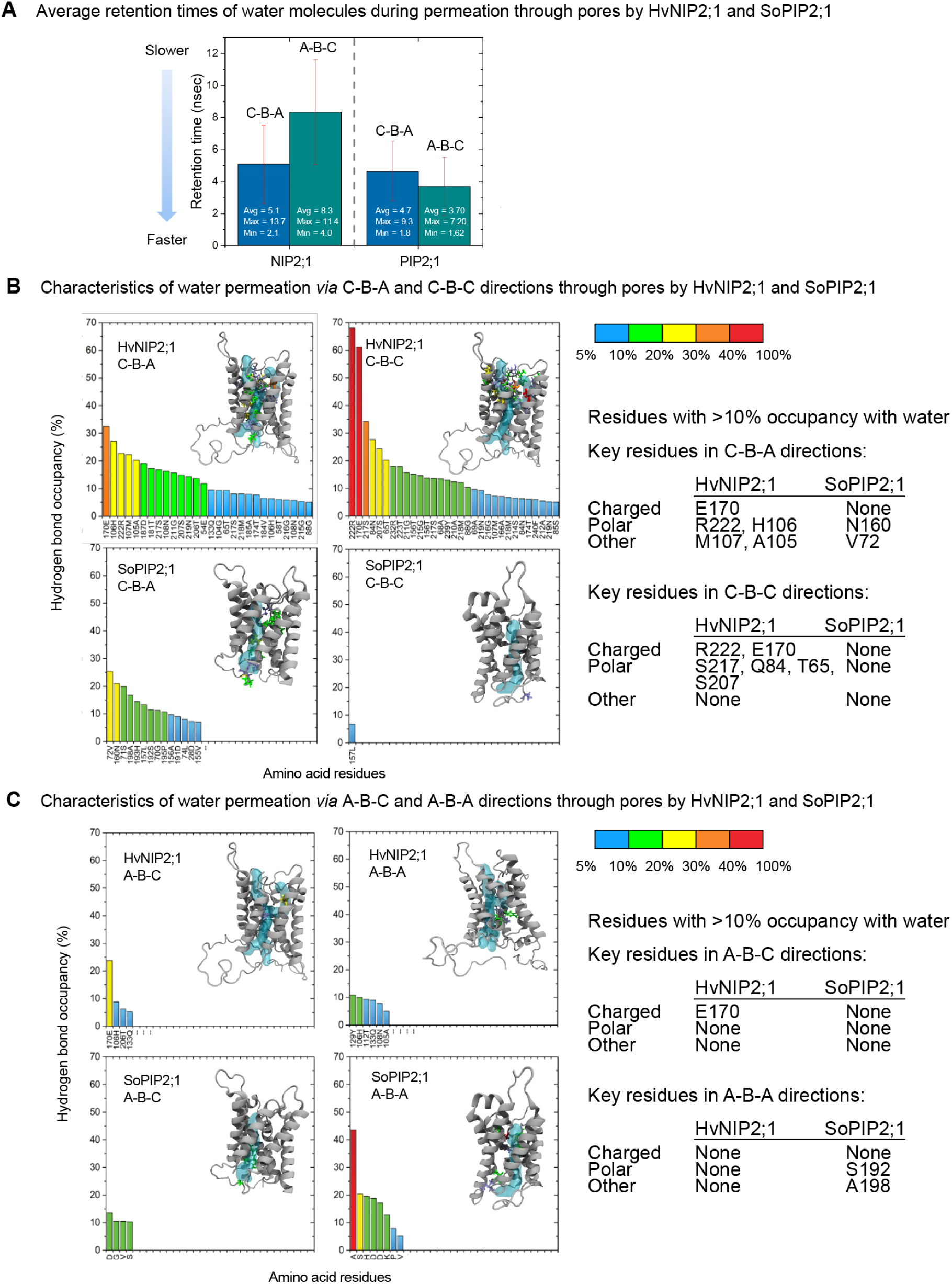
Water permeation characteristics of HvNIP2;1 and SoPIP2;1 based on steered MD simulations. *(A)* Average retention times (nsec) of water molecules required to traverse pores in A-B-C (all-through pass) or C-B-C (U-turn) directions. *(B)* Histograms of hydrogen bond occupancy of residues (in %, coloured in blue, green, yellow, and red) with water molecules for preferred C-B-A and C-B-C directions. Insets show structures of AQPs with pores as cyan tubes, and with key residues indicated for each AQP. Sticks in cartoons are in colours that correspond to those in histograms, which bind water molecules with occupancy higher than 10%. Key residues involved in binding water molecules are indicated. *(C)* Histograms of hydrogen bond occupancy of residues (in % coloured in blue, green, yellow, and red) with water molecules for A-B-C and A-B-A directions. Descriptions of insets are as those in panel B. Key residues involved in binding water molecules are indicated.

### (i) Water permeation

AQPs (54), including HvNIP2;1 (9) and SoPIP2;1 (38, 39) transport water, and thus molecular mechanisms that govern water transport and permeation directionality were examined through steered MD simulations (Figs. 4 and 10, Fig. S6). The total number of water molecules permeating (passing) through pores per nsec unit time indicated that the C-B-A path was the preferred route for both AQPs, where 23 and 14 molecules passed *via* C-B-A *versus* 8 and 6 molecules *via* A-B-C paths for HvNIP2;1 and SoPIP2;1, respectively (Fig. 4A; Table S3; Movies S1-S8). This quantitative analysis revealed water rectification for the C-B-A paths of each AQP (*i.e.* equivalent to endosmosis), which is a novel observation that now needs to be examined experimentally. Although, in HvNIP2;1 the presence of two C-entrances (Fig. S5A, *cf*. red dashed circles; Movies S1-S4) allowed water molecules to adopt U-turn behavior *via* the C-B-C path, thereby decreasing the overall all-through passing rate. On the other hand, water molecules entering through the A-entrance of HvNIP2;1 made a U-turn at a notably lower rate. Conversely, SoPIP2;1 exhibited an opposite trend (Table S3), where water molecules passing through A-entrance showed a markedly higher U-turn rate than those entering through C-entrance; specifically, 24 and 8 respective water molecules made the U-turn through A-and C-entrances (Table S3). Overall, in SoPIP2;1 39% of all water molecules that traversed the pore had a higher average passing rate through the pore than those in HvNIP2;1 (27%) (Table S3; Movies S1-S8). In addition to water passing directionality, it was important to quantify the simulated speed of water permeation (Fig. 4A) in response to the pulling force, specified above and in Methods. It was envisaged that on average it took approximately the same time for a water molecule to travel through HvNIP2;1 (5.1 nsec) or SoPIP2;1 (4.7 nsec) *via* the C-B-A path, while the A-B-C path showed a higher passing time for HvNIP2;1, making it a less preferred route (Fig. 4A).

To complement the water permeation characteristics of both aquaporins described above with structural analyses, amino acid residues participating in hydrogen bonds with water molecules as they entered pores were identified (Figs. 4B and 4C). These contacts were expressed as occupancies (or time intervals during the last 20 nsec simulations) that were required to form stable hydrogen bonds with water molecules (100% corresponded to a stable hydrogen bond). Data in Fig. 4B indicated that for C-B-A (all-through) and C-B-C (U-turn) routes the number of residues forming strong hydrogen bonds was significantly higher for HvNIP2;1 than for SoPIP2;1. On rare occasions, so-called deep A-B-C to C-B-A U-turns were observed in both AQPs, where water molecules traversed from A-entrance to C-entrance, but didn’t appear to exit at C-entrance, but turned around to exit *via* A-entrance. During water permeation, only pore residues contributed to hydrogen bonds with relatively low occupancy for both AQPs, except R222 (located at a re-entrant α-helix in the selectivity filter) and E170 (at the 5^h^ α-helix facing the central pore) in the C-B-C path for HvNIP2;1 (Fig. 4B; Fig. S10A; Movie S3), and mainly these and H106, S207, S217, Q84, and T65 polar residues (positioned alongside the pore) formed hydrogen bonds with water molecules (Figs. 4B and 4C; Fig. S10A). It was also observed that in SoPIP2;1 A198 and S192 (which are relatively small) residues contributed to a high U-turn in the A-B-A path (Fig. 4C), and that not all residues formed strong hydrogen bonds with water molecules in C-B-A, C-B-C and A-B-C paths in HvNIP2;1, except R222 and E170 as previously stated (Figs. 4B and 4C; Fig. S10A). In SoPIP2;1 through C-B-A (Fig. S10G) and A-B-C routes, only a few residues formed hydrogen bonds with water molecules when located near the pore entries, while after they passed the middle segments of pores, these water molecules adopted an ordered single-file type of arrangements and movements (Movies S5-S8).

### (ii) BA permeation

In HvNIP2;1, contrary to water permeation directionality, the opposite A-B-C path (from the cytoplasm to apoplast) was the preferred route for BA (Fig. 5; Fig. S10B), where this hydroxylated metalloid moved through the pore in a solvated form with tightly bound water molecules. Assuming that in HvNIP2;1 this was the primary permeation route for BA, the strength of interactions between pore residues and BA molecules (Fig. 5A) was examined, where the outer boundary of the pore through which these molecules were travelling, was selected as a reference point (Fig. 5A). It was found that behaviors of HvNIP2;1 and SoPIP2;1 differed, where BA was bound strongly by SoPIP2;1 after it entered the pore, meaning that BA transport through the A-B-C path (Fig. 5A) or C-B-A directions become unviable. Conversely, HvNIP2;1 formed weak non-bonding van der Waals and electrostatic interactions with BA as it travelled in a non-fixed orientation through the pore in the preferred A-B-C direction (Fig. 5A; Fig. S10B). During these processes through the C-B-A path, BA showed stronger interactions in HvNIP2;1, which thus was a less favorable direction for BA passage (Fig. 5A). These molecular motions are shown in representative MD simulation movies (Movies S9 and S10), where each simulation indicates how water molecules locate to the proximity of BA (within 3Å radius). It was found that on average HvNIP2;1 attracted six water molecules solvating BA compared to four water molecules in SoPIP2;1 (Movies S11 and S12). The higher BA solvation occupancy in HvNIP2;1 could reduce the strength of non-bonding interactions between BA and pore residues, and explains the higher mobility of BA in HvNIP2;1, while the lower BA solvation occupancy in SoPIP2;1 made BA permeation virtually impossible (Fig. 5A). Considering the favorable permeation of BA by HvNIP2;1, the structural differences of HvNIP2;1 were evaluated after the BA permeation cycle was completed (Fig. 5B). While no significant differences were detected in HvNIP2;1 for A-B-C or C-B-A paths, small variations were observed near its N-and C-termini (Fig. 5B), which pointed to the overall high stability of these routes. For the A-B-C permeation path in HvNIP2;1, Asn108 and Pro220 were identified as the key residues to form hydrogen bonds with BA together with up to six water molecules, depending on the position of BA in the pore (Fig. S10B).

**Fig. 5.**
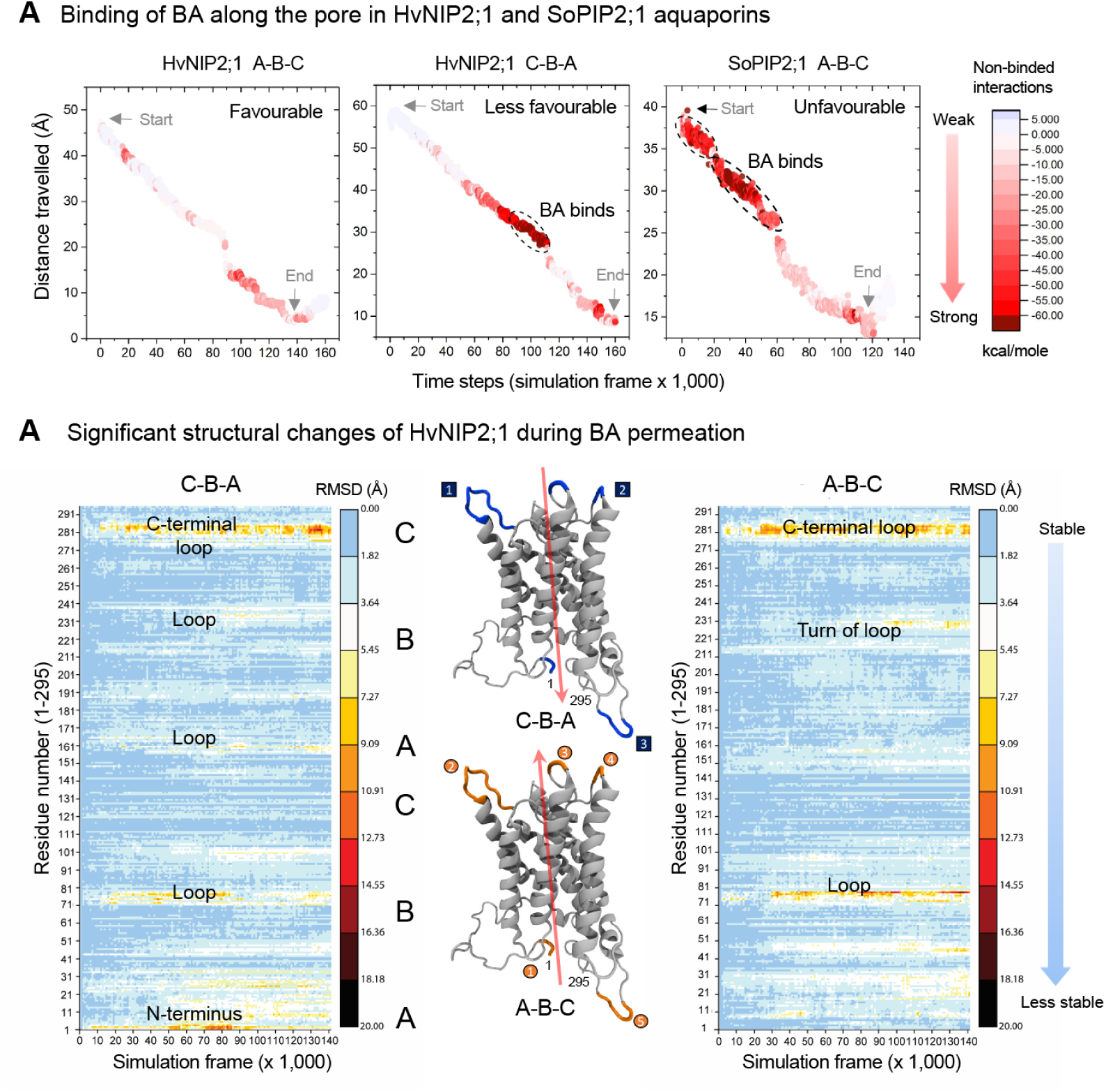
BA permeation characteristics of HvNIP2;1 and SoPIP2;1 based on steered MD simulations. *(A)* Path-depended heat-maps illustrating the strength of non-bonding interactions of BA in pores during permeation in A-B-C and C-B-A (HvNIP2;1) and A-B-C (SoPIP2;1) directions. Dark red circles indicate regions with strong interactions between BA and residues in pores, that preclude BA from migrating through pores (dashed ovals). *(B)* Structural changes in HvNIP2;1 during BA permeation *via* C-B-A and A-B-C directions. Heat maps illustrate changes in residual steered MD values of superposed structures before and after permeation. Blue (C-B-A) or orange (A-B-C) colours in cartoon representations denote regions with high structural changes and those correspond to orange areas in heat maps, while grey colour denotes regions with low structural changes that correspond to white and blue areas in heat maps.

### (iii) Sucrose permeation

Steered MD simulations examining the details of sucrose movements were based on the same approach as described above. Here, the more preferred C-B-A transport path for HvNIP2;1 (from apoplastic to cytoplasmic) for sucrose transport is shown (Fig. 6; Fig. S10C), although it was observed that HvNIP2;1 permeated sucrose through the pore in both C-B-A (Movie S13) and A-B-C (Movie S15) directions. Fig. 6A and Fig. S10C illustrate the spatial dispositions of E170, N108, and T206 residues in the pore of HvNIP2;1 *via* the C-B-A path. These residues formed hydrogen bonds with sucrose, where specifically polar E170 served as an attracting force to bind and retain the disaccharide in the pore; a similar function for E170 was observed in HvNIP2;1 during water permeation. Conversely, SoPIP2;1 experienced a sucrose hold-up in the middle of the pore in both directions (Fig. 6; Fig. S10H-I), consequently, preventing it from moving along the pore (Movie S14 shown for C-B to B-C directions). This hold-up could obviously impose a strong destructive force on SoPIP2;1, where it may lose structural integrity (Fig. 6B; Fig. S10H-right panel, *cf.* white, grey and cyan cartoons). To compare movements of sucrose in HvNIP2;1 and SoPIP2;1, temporal profiles of time-depended non-bonding van der Waals and electrostatic interactions that were formed between pore residues and sucrose, were investigated during the 80 nsec MD simulations run (Fig. 6B). In HvNIP2;1, these non-bonding interactions representing mainly hydrogen bonds, were key contributors to sucrose movements during entire runs, while in SoPIP2;1, although comparable at the beginning, these interactions dissipated mid-way through MD simulations (Fig. 6B, right panel). Changes in structural regions of HvNIP2;1, when the pulling force as specified above was imposed, were identified with high (orange) and low (grey) impacts, while heat maps illustrated average displacements along the pore compared to the original HvNIP2;1 state (Fig. 6C). When comparing sucrose permeation by HvNIP2;1 in both directions, it became obvious that the preferred route for HvNIP2;1 was the C-B-A path, and as for this path, HvNIP2;1 maintained high structural integrity (excluding N-and C-termini that are expected to be mobile). Conversely, the less preferred A-B-C path led to structural disruptions (Fig. 6C). In a biological context, given that the N-terminus of HvNIP2;1 is cytoplasm-oriented (42, 43), the preferred C-B-A sucrose permeation path, revealed *via* steered MD simulations, implies that sucrose could be transported from extracellular to cellular environments, and less so in the opposite direction.

**Fig. 6.**
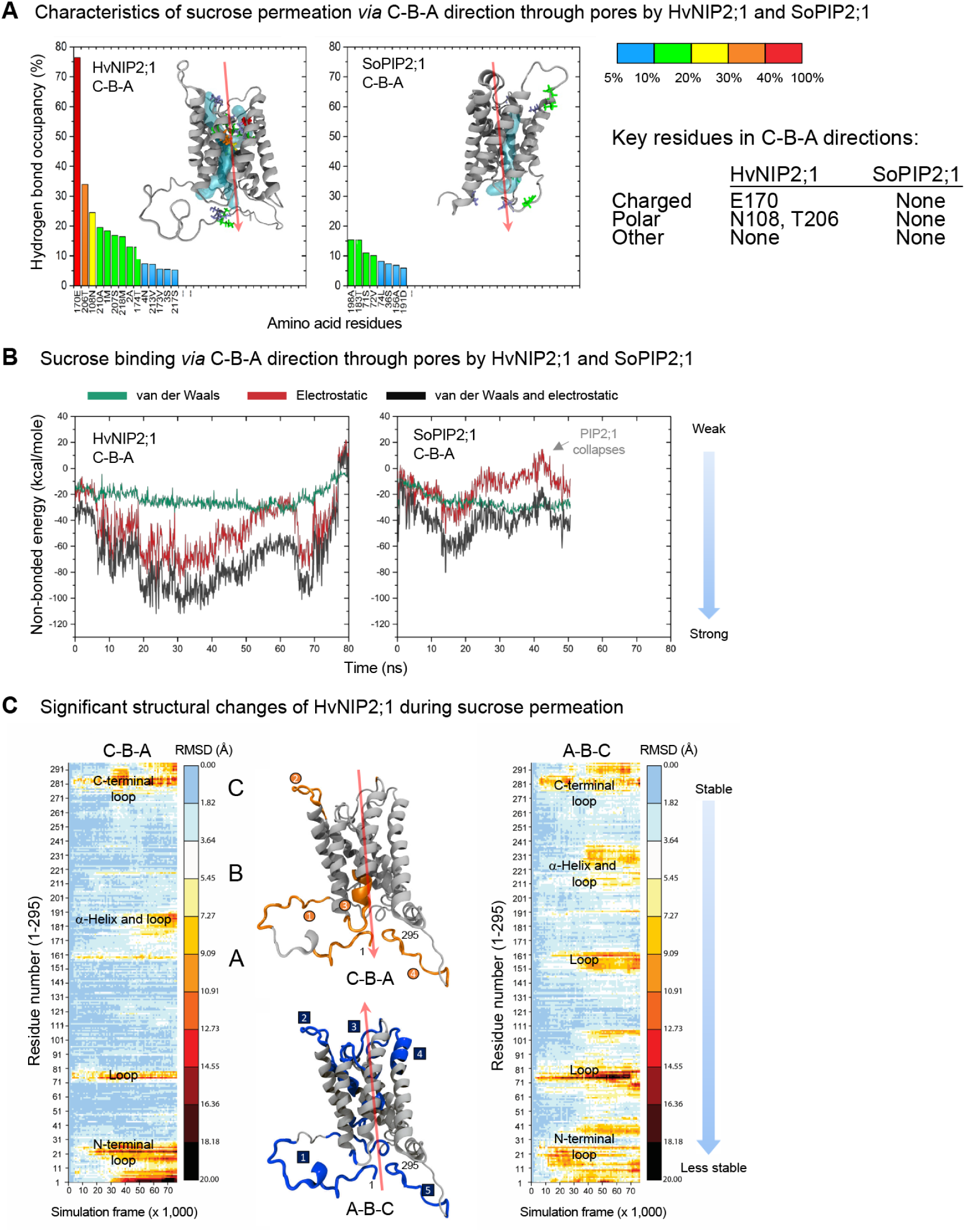
Sucrose permeation characteristics of HvNIP2;1 and SoPIP2;1 based on steered MD simulations. *(A)* Histograms of hydrogen bond occupancy of residues (in %, coloured in blue, green, yellow, and red) with sucrose for the C-B-A direction. Insets show structures of AQPs with morphologies of pores (cyan tubes), and key residues that bind sucrose with occupancy higher than 10%. *(B)* Time-depended quantification of molecular forces involved in binding sucrose. Calculations of van der Waals (green trace), electrostatic (red), and the sum of both forces (total energy; blue) are indicated for the preferred C-B-A direction for both AQPs. While HvNIP2;1 permeates sucrose, SoPIP2;1 fails and its structure collapses midway through permeation. *(C)* Structural changes of HvNIP2;1 during sucrose permeation *via* C-B-A and A-B-C directions. Heat maps illustrate changes in residual steered MD values of superposed structures before and after permeation. Blue (C-B-A) or orange (A-B-C) colours in cartoon representations denote regions with high structural changes and those correspond to black, yellow, and red areas in heat maps, while grey colour denotes regions with low structural changes that correspond to white and blue areas in heat maps.

**Fig. 7.**
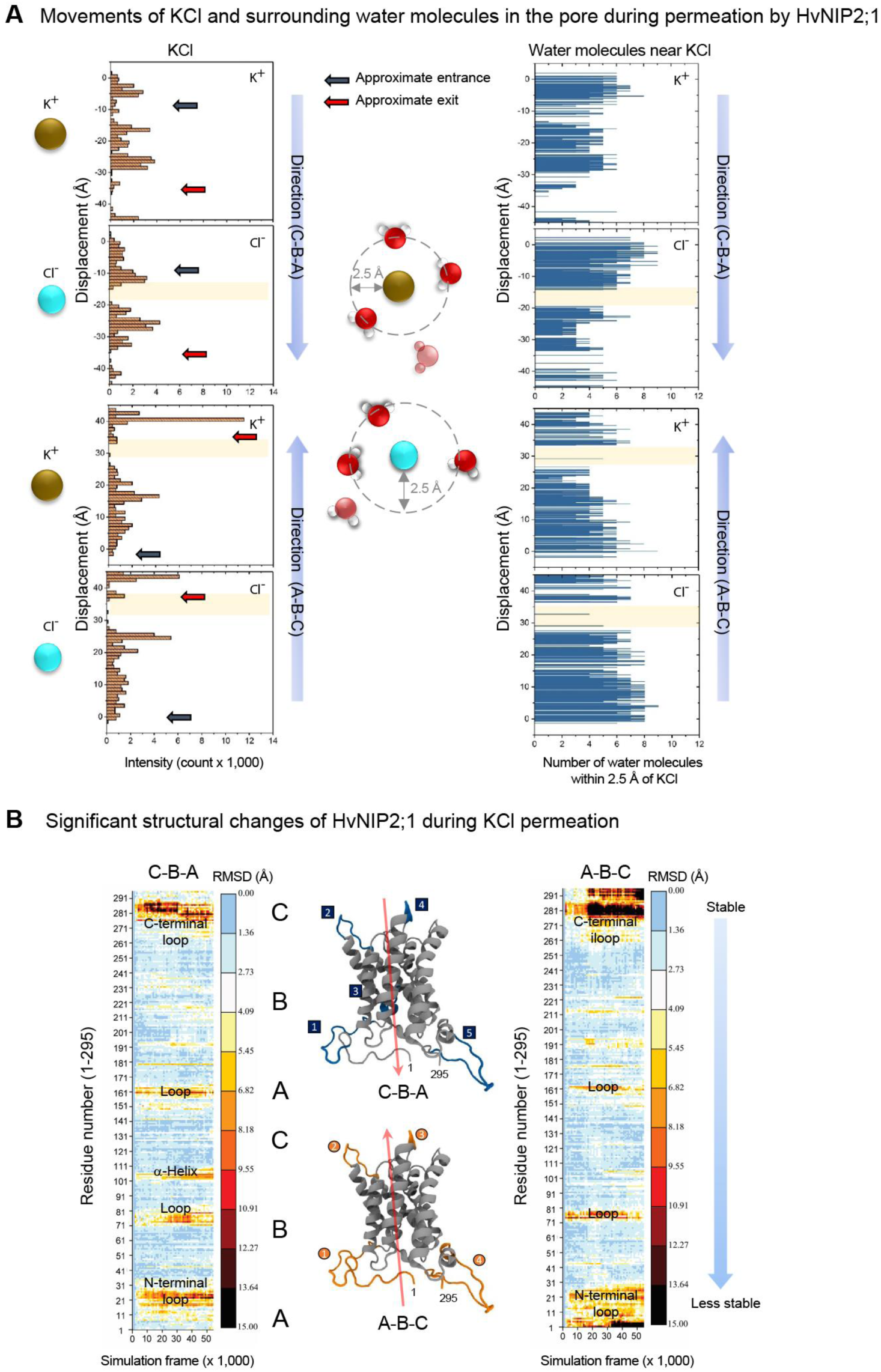
KCl permeation characteristics of HvNIP2;1 based on steered MD simulations. *(A)* Left panels: Trace maps for K^+^ and Cl^-^ ion pairs that travel through pores and interact with pore residues *via* C-B-A and A-B-C directions. Intensity correlates with the likelihood of an ion pair staying in a particular location in a pore, and displacement denotes the path of ion pairs from starting points. Regions with sudden motions of ions through pores (not accompanied by water molecules) are highlighted in light orange. Right panels: Trace maps of water molecules (at a distance of ≤2.5 Å) surrounding K^+^ and Cl^-^ ion pairs that are transported through pores. Regions lacking surrounding water molecules are highlighted and match those in left panels. *(B)* Structural changes of HvNIP2;1 during KCl permeation *via* C-B-A and A-B-C directions. Heat maps illustrate changes in residual steered MD values of superposed structures before and after permeation. Blue (C-B-A) or orange (A-B-C) colours in cartoon representations denote regions with high structural changes and those correspond to black, yellow, and red areas in heat maps, while grey colour denotes regions with low structural changes that correspond to white and blue areas in heat maps.

### (iv) KCl permeation

Given that KCl was shown experimentally to permeate relatively rapidly through HvNIP2;1 (Fig. 2; Fig. S1, Table S1), MD simulations of KCl transport were used to quantify the motions of K^+^ and Cl^-^ ion pairs carried out through A-B-C and C-B-A directions in both AQPs (Fig. 7; Figs. S7; S10D-E, S10J-K; Movies S17-S20). Representative images of K^+^ and Cl^-^ ion pairs, and associated water molecule movements indicated the positions of both ion pairs along NPA1-and NPA2-motifs and other residues in the vicinity of ion pairs in HvNIP2;1 (Fig. S10D-E) and SoPIP2;1 (Fig. S10J-K). Approximate entrances (blue arrows) and exits (red arrows) points (Fig. 7A; left panel) were identified along with regions where sudden ion pair motions occurred; these sudden motions need to be investigated further. When examining the permeation of K^+^ and Cl^-^ ion pairs, these were considered as being solvated in water, and thus simulations were carried out using hydrated states at a distance less or equal to 2.5 Å from ions. Fig. 7A (left panels, highlighted in light orange) illustrates the movements of K^+^ and Cl^-^ ions through the pores of HvNIP2;1, although in some pore regions, these ions were not accompanied by water molecules (Fig. 7A, right panels; highlighted in light orange). These regions, where ions lacked accompanying water molecules, matched those regions where ions in both AQPs became immobilised (Figs. 7A; Fig. S7A, *cf.* left and right panels), implying that this could be the explanation for these ion hold-ups. Hence, it could be concluded that ion pairs had difficulties during the traversing of pores of both AQPs without associated water molecules (Figs. S10D-E; S10J-K). It was further observed for HvNIP2;1 (Fig. 7; Movies S17 and S18) and SoPIP2;1 (Fig. S7; Movies S19 and S20) that K^+^ and Cl^-^ ion pairs were permeated in both A-B-C and C-B-A directions (Fig. S10D-E). During KCl ion pair permeation in HvNIP2;1, G104, A105, and P109 appeared to participate in van der Waals interactions, contingent on the positions of ions in the pore (Figs. S10D and S10E, shown for A-B-C and C-B-A directions, respectively). Structural alterations were examined in both AQPs before and after KCl permeation (Fig. 7B; Fig. S7B). For HvNIP2;1, significant changes in the structure would occur in both directions (Fig. 7B), while these changes were mainly observed outside of the pore at the edges of HvNIP2;1 and near N-and C-termini (Fig. 7B). A somewhat similar situation was observed for SoPIP2;1, although in this case, the extent of structural changes was higher (Fig. S7B) compared to that of HvNIP2;1, and these structural perturbations occurred at several α-helices near entrances to the pore (Fig. S7B, blue 1-4 and orange 1-5 marks for C-B-A and A-B-C directions). Based on steered MD simulations with the applied pulling force it could be suggested that both AQPs could permeate KCl in both directions.

### (v) NaNO_3_ permeation

Steered MD simulations revelated the extent of motions of Na^+^ and NO3**^-^** ion pairs alongside the C-B-A and A-B-C directions of HvNIP2;1 and SoPIP2;1 (Figs. S8, S9, S10F, S10L; Movies S21-S24). Fig. S8 shows abnormalities in the Na^+^ and NO3**^-^** ion pair permeation through the A-B-C path in HvNIP2;1. When this ion pair enters the pore, the Na_+_ component of the ion pair freezes for a significant time, and then it suddenly moves down the pore under the force of steered MD simulation (Movie S21). Since this event did not occur during the C-B-A path, such behavior subtly suggested that the support for NaNO_3_ permeation in the A-B-C direction in HvNIP2;1 is weak; these MD simulation data matched our experimental observations of weak NaNO_3_ transport rates by HvNIP2;1 (Fig. S1). As for accompanying water molecules of ion pairs during the A-B-C permeation route, no irregularities were observed at distances less or equal to 2.5 Å to Na^+^ and NO3**^-^** ions (Fig. S8A; right panels). SoPIP2;1 behaved differently, where several structural regions were identified containing immobilised ions, and where these ions continued to move along the pore in C-B-A and A-B-C directions (Fig. S9A, highlighted in light orange; Movies S21 and S22). Fig. S9A visualises the pore areas in SoPIP2;1, where Na^+^ and NO_3**-**_ ions lacked accompanying water molecules and which again matched ion immobilisation areas. These observations emphasised the significance of accompanying water molecules for uninterrupted NaNO_3_ permeation. Analyses of residues involved in NaNO_3_ permeation through the middle section of the HvNIP2;1 pore in the C-B-A direction indicated that the NPA2 selectivity filter residues Asn219 and Pro220 formed van der Waals interactions with ion pairs (Fig. S10F), and that structural changes observed in the pore during NaNO_3_ permeation (Fig. S8B) were similar to those of KCl (Fig. S7B). When comparing the A-B-C and C-B-A paths for NaNO_3_ permeation in HvNIP2;1, it was noticed that during the C-B-A path, the structure altered less than during the A-B-C path, which could make the former path a preferred route. On the other hand, harsh structural perturbations were observed in SoPIP2;1 during NaNO_3_ permeation *via* the C-B-A path, while during the opposite A-B-C path only α-helices at the cytoplasmic side of SoPIP2;1 were affected (Fig. S9B). Considering this behavior and analyses of steered MD simulations (Movies S23 and S24), it was concluded that permeation of NaNO_3_ in SoPIP2;1 *via* both routes was less likely to occur or that this ion pair was permeated at a low rate.

### Phylogenomic analyses of MIP proteins

Randomised Axelerated Maximum Likelihood (RAxML) phylogenetic analysis of Chlamydomonas reinhardtii, Volvox carteri, Physcomitrella patens, Amborella trichopoda, Spirodela polyrhiza, Arabidopsis thaliana, and Hordeum vulgare MIP sequences were used to define phylogenetic relationships and variations in selectivity filter residues of MIPs (Fig. 8; Fig. S11). These analyses indicated that 164 Viridiplantae (Fig. 8) and additional 2,993 archaean, bacterial, fungal, and metazoan entries of the MIP family (Pfam database PF00230) (Fig. S11; Dataset S1) clustered in four major TIP, NIP and SIP clades, where PIP and TIP clades covered the majority of MIP entries. Barley HvNIP2;1 (Fig. 8, bold) with the GSGR selectivity filter residue signature, was resolved to form a monophyletic group with Arabidopsis thaliana and Amborella trichopoda, and that HvNIP2;1 split from Physcomitrella patens that carried the FAAR signature. The three NIP1, NIP2, and NIP3 clades emerged prior to the evolution of tracheophytes (ferns and seed plants) with NIP2 and NIP3 clades containing basal Physcomitrella sequences, and with the NIP1 representative possibly lost. The functional evolution of NIP clades was evident, where the selectivity filter in Physcomitrella (FAAR) diversified into the GSGR signature in the NIP3 sub-clade, following the tracheophyte/Physcomitrella split. This contrasted with PIP entries, where the FHTR signature was conserved (Fig. 8). The GSGR selectivity filter residue pattern in NIP3 entries supports the permeation of larger solutes such as hydroxylated metalloids and certain saccharides (Figs. 2, 3, 6; Figs. S1-S2; Table S1). Conversely, in PIPs, TIPs, and SIPs these signatures contained bulky residues (F for PIPs; His, N, Q, M or I for TIPs; W, F, Y, V or T for SIPs), but also W for NIP1 and V, A or T for the NIP2 clade entries in the 1_st_ positions, compared to NIP3 entries (Fig. 8). These residue characteristics agreed with published data (2, 33, 56–58), where HvNIP2;1 consistently clustered in the NIP3 sub-clade.

**Fig. 8.**
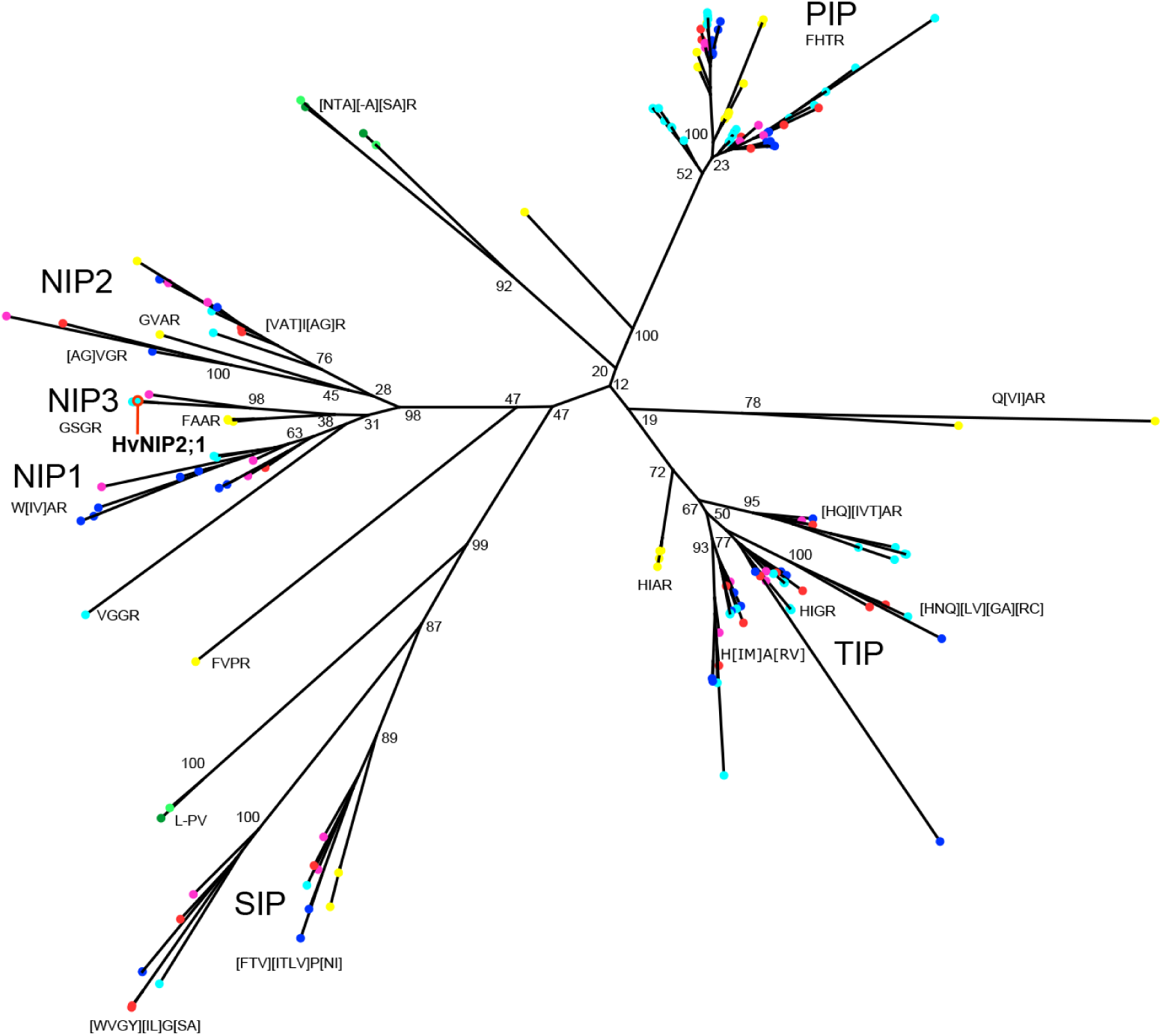
Evolutionary relationships of Viridiplantae MIP proteins. RAxML tree of 164 Viridiplantae MIP proteins. Terminal nodes are colour coded by species: dark green, *Chlamydomonas reinhardtii*; light green, *Volvox carteri*; yellow, *Physcomitrella patens*; magenta, *Amborella trichopoda*; red, *Spirodela polyrhiza*; cyan, *Hordeum vulgare*; blue, *Arabidopsis thaliana*. A red circle marks HvNIP2;1 (in bold) investigated in this work. Selectivity filter residues are noted adjacent to relevant clades and residues enclosed in square brackets indicate variations at those positions. Bootstrap support values are indicated at major nodes.

## Discussion

In the present work, we observed that HvNIP2;1 when reconstituted in liposomes, permeated large cyclic saccharide molecules such as sucrose, L-arabinofuranose, and lactose but not D-glucose of D-fructose (Fig. 2; Figs. S1 and S2; Table S1). This is a novel observation not reported previously for any AQP, including other structurally similar NIPs or NOD26. AQP9 from humans and a rat were found to transport large molecules including polyols (glycerol, D-mannitol, D-sorbitol), purines (adenine), pyrimidines (uracil, 5-fluorouracil), and thiourea, but not cyclic saccharides (59, 60). To define the saccharide specificity permeation by HvNIP2;1, an extensive panel of substrates was examined. Here, neutral (D-xylopyranose, D-glucopyranose, D-fructopyranose, D-galactopyranose, D-mannopyranose and D-mannopyranopheptaose) and charged (D-glucosamine, N-acetyl β-D-glucosamine, and D-glucuronic acid) monosaccharides, disaccharides (trehalose, cellobiose, gentiobiose), and the trisaccharide raffinose were permeated at lower rates than sucrose, L-arabinofuranose and lactose (Fig. 2; Figs. S1 and S2; Table S1).

The next group of permeants transported by HvNIP2;1 included KCl and MgSO_4_ ion pairs, while others such as CH_3_COONa and NaNO_3_ were permeated at lower rates (Fig. 2; Fig. S2) and NaF was impermeable (Fig. S2). These observations support previously observed electrogenic ion conductance through NOD26 (structurally similar to HvNIP2;1) when embedded in lipid bilayers (26). Although, our observations that HvNIP2;1 embedded in liposomes permeated both KCl and MgSO_4_ ion pairs, is a new observation for any NIP-type AQP. It remains to be established if this transport also translates to an electrogenic ion conductance as opposed to neutral ion pair permeation. Notably, previous studies described electrogenic ion conductance in human AQP1 and AQP6, AtPIP2;1 (that has different features compared to SoPIP2;1) and AtPIP2;2, HvPIP2;8 and OsPIP1;3, when expressed in oocytes or HEK 293 cells (61–65). It was theorized that electrogenic ion conductance could proceed through a central pore, where four monomeric AQPs converge, and that at least in human AQP1 this function could be important for cGMP-mediated gating (36). Nevertheless, neutral ion pair permeation in some AQPs could occur *via* the monomer pores as indicated by steered MD simulations shown here.

HvNIP2;1 embedded in liposomes, also permeated hydroxylated metalloids BA and germanic acid and some other neutral solutes known to be permeated by AQPs (Figs. S1-S2; Table S1). Some permeability to BA, urea, and glycerol was observed in control liposomes lacking HvNIP2;1 or in proteo-liposomes incubated with AgNO_3_ (Fig. 2C), as these small molecules could permeate passively through lipid bilayers (50, 66). It was observed that the osmotic water permeability coefficient of HvNIP2;1 (*P*_water_ = 3.98 ×10^-2^ cm s^−1^) was 240-fold higher than that of empty liposomes (Table S1) and similar to the liposome-reconstituted *Escherichia coli* water-permeable aquaporin-Z (*P*_water_ = 2.83 ×10^-2^ cm s^-1^) (67) but higher than that of NOD26 (*P*_water_ = 1.2 ×10^-2^ cm s^-1^) (68). The comparison of *P_BA_* = 2.50×10^-6^ cm s^-1^ for HvNIP2;1 with *P_BA_* of tapetal NIP7;1 (1.41×10^-6^ cm s^−1^) involved in pollen cell wall formation (69), indicated that these values were similar, although in these analyses it is imperative to consider differences in the protein content of liposomes, embedding directionality and protein homogeneity. As for other solutes such as urea, it was surprising but not unexpected that in HvNIP2;1, its permeation rate compared to that for BA, differed (Fig. S1; Table S1), although their molecular radii are similar (BA - 2.56 Å; urea - 2.30 Å), which agreed with data of rice Lsi1 (70) and soybean NOD26 (71), where low or no urea permeability was reported.

These findings were validated through oocyte swelling assays with HvNIP2;1 expressed in *X. laevis,* where water, BA, and sucrose transport but not glucose transport occurred in the CBA direction (into oocytes) (Fig. 3), confirming stopped-flow recordings with HvNIP2;1 in proteo-liposomes, whereas after the application of AgNO_3_ permeability was completely suppressed. Notably, BA and sucrose did not compete during transport when utilised together suggesting that they could be co-permeated (Fig. S3).

A comprehensive set of steered MD simulations revealed how HvNIP2;1 and SoPIP2;1 AQPs permeated solutes that for HvNIP2;1 were examined experimentally in this work. These simulations, while pulling ligands through pores *via* an external force, reveal details of ligand binding and energy landscapes, and probe conformational changes, ligand-residue interactions, and mechanical and elastic properties of proteins at nsec scales (53). These parameters cannot often be accessed experimentally (53). Naturally, the outcome of the steered MD simulations will rely upon the design of the system, and on the careful choice of simulation parameters. Specifically, steered MD simulations require the optimisation of direction, and strength of the force *versus* velocity of ligand pulling, to avoid structural collapse. One of the advantages of steered MD simulations over conventional ones is to induce protein conformational changes on nsec scales (53).

Simulating the permeation events using steered MD simulations with monomeric HvNIP2;1 and SoPIP2;1 provided snapshots of how solutes pass through pores. In the future, these studies could be extended to tetramers with higher computational costs (72). In our simulations, we used optimised parameters that will play a key role in generating realistic results, where the derivation of these parameters requires trial-and-error runs (73). Following the logistics from our previous works (74), the best combinations of force (kcal/mol) and velocity (Å/time step) values were selected, where a spring constant of 0.1 kcal/mol/Å_2_ and a pull rate of 0.05 fN per a 2 fsec simulation time-step or a vector heading from the entrance of the pore to the opposite side of the pore. Using this pulling force, it was confirmed for the HvNIP2;1 model and the SoPIP2;1 crystal structure that these AQPs exhibited water transporting characteristics (Fig. 4) when examining A-B-C (cellular to extracellular) and C-B-A (from extracellular to cellular environments) routes. Both AQPs preferred the C-B-A direction for water permeation (endosmosis in a cellular context) and in HvNIP2;1 E170, S217, and R222 residues assisted and sourced crucial nonbonding van der Waals interactions (Fig. 4B). Conversely, the more preferred permeation route for BA by HvNIP2;1 was the A-B-C path and this was due to the concomitant presence of a significant number of water molecules that solvated BA. These water hydration shells reduced potentially strong interactions of BA with residues at the pore entrance through the A-B-C path (Fig. 5A; Fig. S10A). There were no significant structural differences in HvNIP2;1 during BA transport *via* A-B-C or C-B-A routes (Fig. 5B), with hydrogen bonds formed between BA and E170, N108, and M218 (Fig. 5A; Fig. S10B).

Sucrose permeation by HvNIP2;1 was also guided by both electrostatic and van der Waals interactions (Fig. 6), where HvNIP2;1 contributed with E170, N108, and T206 as pivotal residues in the preferred C-B-A direction (Fig. 6; Fig. S10C). To explain the inability of HvNIP2;1 to permeate smaller monomeric D-glucose or D-fructose molecules compared to the disaccharide sucrose (a larger molecule) that we observed, the schematics of residue distributions (Fig. S5) informs that the voluminous HvNIP2;1 pore is decorated with hydrophobic and hydrophilic residues at a ratio of 17:18 (with two negatively and one positively charged residues) (Figs. S5 and S12). We assume that in the HvNIP2;1 pore, hydrophilic glucose could get trapped in pockets, while sucrose, due to its larger size avoids this trapping and slides through the pore. Conversely, a narrower and less capacious SoPIP2;1 pore (Fig. S5) and the hydrophobic/hydrophilic residue ratio 19:6 (Fig. S12) would collectively offer an unfavorable milieu for saccharide molecules to pass through.

In planta, HvNIP2;1 AQP is expressed in roots and localised to epidermal and cortical cells of seminal roots and hypodermal cells in lateral roots (9, 10). The biological significance of our key findings of sucrose and ion permeation means that HvNIP2;1 could carry these permeants along with water and metalloids and that this could have profound importance in plant physiology (35, 75). Through this permeation pathway, HvNIP2;1 located on root cell membranes could recover sucrose through the preferred C-B-A path directed from apoplastic (extracellular) to cellular environments. Consequently, this permeation route could constitute the novel element of a plant’s cellular saccharide-transporting machinery.

The support for our original *in-vitro* and *in-silico* findings that HvNIP2;1 conducted KCl and to a lesser extent NaNO_3_, is in structural differences of pores, but it was also noticed that these differences between A-B-C and C-B-A directions were subtle (Fig. 7; Figs.S7-S9), while differences near pore entrances were larger. It is reasonable to suggest concentrations of such ion pairs in biological solutions are in μM ranges. Suggest it will be in micro molar range. It was observed that as solvated K_+_ and Cl_-_ ion pairs moved along the pore of HvNIP2;1, in the vicinity of NPA1-and NPA2-motifs, these residues assisted in binding solvated ion pairs that did not shed water molecules from their shells (Fig. S10D-E). This shedding would obviously create a large energy cost to permeation. It was also noted that HvNIP2;1 showed a preference for the C-B-A path for solvated Na^+^ and NO_3-_ ion pairs (Fig. S10E), while SoPIP2;1 did not permeate this ion pair.

When using steered MD simulations, it is important to relate the external time-dependent force to parameters typically used in *in vitro* or *in vivo* experiments. It was noted that the force applied in simulations can be related to the hydrostatic pressure difference between the two sides of a membrane that produces a net directional water flow (76). But this difference cannot be implemented under periodic boundary conditions, as biological membranes are bound by water layers on each side. To implement realistic force parameters, cautious approaches are needed when applying temperature, pH, and ionic strength constraints, which could easily be adjusted to predict, *e.g.* permeability coefficients inaccessible experimentally (72, 77–79).

To investigate the evolutionary origin of NIP AQPs, the phylogenetic reconstruction of 164 Viridiplantae MIPs was conducted to reveal that PIP, TIP, and SIP clades diversified before the chlorophyte (green algae) and embryophyte (land plants) split, proposing that a single duplication event resulted in SIP, or that multiple duplication events occurred before the chlorophyte and embryophyte split. These analyses use a relatively complex substitution model avoiding insufficient data for maximum likelihood but still keeping accuracy, although there was little support for establishing deep relationships in MIPs as these sequences are relatively short. Hence, we restricted maximum likelihood analyses to Viridiplantae only (Fig. 8), and separately to Virdiplantae, archaean, bacterial, fungal, and metazoan entries (Fig. S11; Dataset S1). While we observed an increased gene duplication in angiosperms (flowering plants) in all analysed clades, our phylogenetic reconstruction showed that there were ancestral gene losses in chlorophytes, although deep relationships between NIP, PIP, and TIP entries were obscured by poor node support (Fig. 8; Fig. S11). Notably, and relevant to the experimental data obtained in this work, the NIP3 sub-clade was segregated clearly and carried the GSSR selectivity filter signature. This NIP3 clade was lost or reduced in eudicots, although the deep phylogenetic relationships of AQP clades were less supported (Fig. S11). Nevertheless, the NIP3 clade segregation suggested that the pore residues were prone to alterations to modulate solute selectivity in a NIP3 sub-clade to allow the permeation of hydroxylated metalloids and larger molecules such as sucrose (Fig. 2; Fig. S1). Meanwhile, PIP entries positioned on short molecular branches, showed strong selectivity filter residue conservation in Viridiplantae, while NIP, TIP, and SIP entries underwent more substitutions over time. In agreement with published data (33, 56–57), our analysis confirmed that the three NIP1-NIP3 sub-clades diversified early during embryophyte evolution, before the split from the tracheophytes. Further, the NIP3 sub-clade separated from those of NIP1 and NIP2 AQPs or appeared to be lost or reduced in eudicots (Fig. S11), and NIP3 entries showed clear dichotomy in GSSG and FAAR selectivity filter residues (Fig. 8), which ultimately dictates solute selectivity. This event may have resulted in gaining selectivity that would allow NIP3 members to permeate specific saccharides, as shown in this work. In conclusion, our phylogenomic data of 3,157 AQPs propose that HvNIP2;1 acquired a unique solute specificity as a saccharide transporter.

## Materials and Methods

Chemicals and procedures used for protein expression, solubilisation, purification, and liposomal reconstitution of HvNIP2;1 are detailed in Supplementary Information.

### Stopped-flow light scattering recordings of solute transport in proteo-liposomes with HvNIP2;1

Permeability of DMPC proteo-liposomes with reconstituted HvNIP2;1 and control liposomes lacking HvNIP2;1 was measured to test the transport of 11 solutes (46) with the DX.17MV stopped-flow spectrophotometer (Applied Photophysics, Leatherhead, UK). The shrinking and re-swelling of vesicles were measured by 90° light scattering (550 nm) at 21 _o_C, upon rapidly mixing solutions that create an outward-directed concentration gradient with test solutions in the liposome buffer [20 mM Tris-HCl (pH 8.0) containing 100 mM KCl] at the concentration of 0.2 M (340 mOsmol/kg). These measurements were repeated and extended, such that in total, the transport of a panel of 27 solutes was investigated. Traces from five individual stopped-flow acquisitions were averaged and normalised shrinking and swelling kinetics were fitted to a non-linear regression single exponential function, from which rate constants were calculated. To inhibit solute permeation, the thiol-group modifier AgNO_3_ at 0.5 mM was used (50). Stopped-flow light scattering data were analysed using Prism 9.0.0 (GraphPad Prism Software, Inc., San Diego, CA, USA) based on two biological and two technical replicates of five averaged stopped-flow acquisitions, using non-linear regression of a one-phase decay (solutes) or two-phase association (water) models. Osmotic permeability *P* coefficients for water were calculated based on *P_water_* = (V/A) × rate constant / (V_w_ × C_o_) (78) and permeability coefficients of all other solutes based on *P_solutes_* = (V/A) × rate constant (80), where V/A is the volume to surface area ratio of liposomes, V_w_ is the partial molar volume of water and C_o_ is the external osmolarity after mixing, using the 50-nm liposome radius. *P_solutes_* coefficients were corrected for nonselective diffusion through lipid bilayers using control liposomes. Diameters of DMPC liposomes with reconstituted HvNIP2;1 and control liposomes were determined using the NICOMP 380 Particle Sizing System (Santa Barbara, CA, USA) operating in a vesicle mode. The data were weighted on ±60 liposomes.

Heterologous expression of HvNIP2;1 in *Xenopus laevis* oocytes and oocyte swelling analyses are described in Supplementary Information.

### Steered MD simulations of BA, sucrose, and KCl and NaNO_3_ ion pair permeation by HvNIP2;1 and SoPIP2;1

Details of the preparation of the full-length monomer HvNIP2;1 model and MD simulations of HvNIP2;1 and SoPIP2;1 are described in Supplementary Information. The 3D molecular model of HvNIP2;1 was constructed based on the structural information available before the coordinates of rice HvNIP2;1 (PDB 7nl4 and 7cjs) were released on 3 November 2021 (42, 43). In these structures, respective 44 and 24(7cjs) or 22(7nl4) N-and C-terminal residues remained unresolved. Steered MD simulations of BA, sucrose, and KCl and NaNO_3_ ion pairs for the HvNIP2;1 model and the SoPIP2;1 crystal structure (PDB 2B5F) (38, 39) were performed using NAMD 2.12 (81), where solute permeation events were examined by pulling solutes through pores from one side of the structure to the other. Solutes were harmonically constrained with *k* spring force constant, to move with *v* velocity, in a given direction, and the vector heading towards the center of the opposite entrance. The potential energy *U* was applied, where *v* is the actual position of the atom, 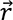 is the initial position of the atom, *t* is time, and 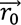 is the direction of pulling:

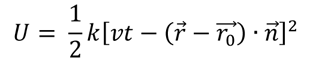

The coordinates of HvNIP2;1 and SoPIP2;1 were taken from the last frame of 100-ns MD simulation trajectory for all steered MD inputs. In accordance with our previous protocols (73), the force (kcal/mol) and velocity (Å/time step) combination parameters were investigated with AQPs, where up to eight steered MD simulations were performed to determine the best combinations. Here, target permeated molecules were harmonically constrained with a spring constant of 0.1 kcal/mol/Å_2_ and a pull rate of 0.05 fN per a simulation time-step (2 fsec) (or 0.025 N/s) in the pore direction, which is a vector heading from the entrance of the pore towards an opposite side of the pore. These spring constant values along with the pull rate matched those established in our previous studies with other systems (82). All bonding and non-bonding parameters including partial charges for sucrose, BA, KCl and NaNO_3_ were obtained from GLYCAM (83) and general AMBER force field (84), respectively. Steered MD simulations were run up to 80 nsec with a time step of 2 fsec. Analyses of root-mean-square-deviation (RMSD) values, and distance and energy parameters were made in the CPPTRAJ module (85) of AMBERtool16 and VMD 1.9.3 (86). HOLE (87) and MOLE 2.0 (88) were used for geometry evaluations of permeation pathways. Lists of residues involved in these pathways of HvNIP2;1 and SoPIP2;1 and phylogenomic analyses are specified in Supplementary Information.

## Acknowledgments

This work was supported by funding from the Australian Research Council (DP120100900) to M.H., and the Australian Research Council Centre of Excellence in Plant Energy Biology (CE1400008) to S.D.T. A.V. was supported by the Sastra University scholarship (Thanjavur, Tamilnadu, India), J.G.S. from the University of Adelaide (Australia), and M.H. from the University of Adelaide and the Huaiyin Normal University (China). Computational work was carried out at the North Carolina State University High-Performance Computing Center and by the Center for Lignocellulose Structure and Formation, an Energy Frontier Research Center (USA) funded by the U.S. Department of Energy, Basic Energy Sciences Award DE-SC0001090.

## Notes

### Competing Interest Statement

The authors have declared no competing interest.

