## Supporting Information for "Barley HvNIP2;1 aquaporin permeates water, metalloids, saccharides, and ion pairs due to structural plasticity and diversification"

**This PDF file includes:**

Supplementary Materials and Methods
Supplementary Results
Figures S1-S12
Tables S1-S3
Legends for Movies S1-S24
Legend for Dataset S1
Supplementary References

**Other supplementary materials for this manuscript include the following:**

Movies S1 to S24
Dataset S1

**Supplementary Materials and Methods**

**Materials.** Oligonucleotide primers, restriction enzymes, plasmid extraction kits, the EDTA-free Complete Protease Inhibitor Cocktail, iohexol (Accudenz), Blue Dextran, and other chemicals were sourced as described (1-4). Benzonase (40 U/ml) was purchased from Invitrogen (Carlsbad, CA, USA), 1,2-dimyristoyl-sn-glycero-3-phosphocholine (DMPC) was from Avanti Polar Lipids (Alabaster, AL, USA), and the styrene-maleic anhydride co-polymer 3:1 (SMA) was provided by the courtesy of Dr. Timothy J. Knowles (University of Birmingham, United Kingdom).

**Cloning of HvNIP2;1 and *Pichia pastoris* clone selection.** Cloning of HvNIP2;1 native cDNA (UniProtKB accession D8V828) in the pPICZ (frame B) (Invitrogen) expression vector yielding the native *HvNIP2;1*-Myc-6xHis-pPICZ-B DNA fusion, destined to be transformed in *P. pastoris*, and *Pichia* clone selection were conducted as described (2-4).

**Barley HvNIP2;1 expression in *Pichia pastoris* cells.** Competent X-33 *P. pastoris* cells (Invitrogen) transformed with the linearised *HvNIP2;1*-Myc-6xHis-pPICZ-B were streaked on the YPD plates (composition defined in the EasySelect™ Pichia Expression Kit Manual) containing 100 µg/mL zeocin (InvivoGen, San Diego, CA, USA) and incubated for two days at 28 ºC (3). Cells from a single colony were inoculated into two mL of liquid BMGY media (composition defined in the EasySelect™ Pichia Expression Kit Manual) in 10 mL conical test tubes. Liquid cultures were grown for two days at 25 °C, transferred to 200 mL of liquid BMMY media (composition defined in the EasySelect™ Pichia Expression Kit Manual), and induced with 1% (v/v) methanol in 2-liter Erlenmeyer flasks for four days at 25 °C maintaining 1% (v/v) methanol under shaking (120 rpm; Multitron INFORS HT, Bottmingen, Switzerland). Cells were harvested by centrifugation (4,500xg, 10 min, ambient temperature), pellets resuspended in 10% (v/v) glycerol, and stored at -80 $^{o}$C.

**Fractionation of *Pichia pastoris* cells using urea/alkali treatment.** The distribution of HvNIP2;1 in *Pichia* cellular fractions was evaluated as follows (5). In brief, cells were lysed in the 90% (v/v) Yeast-Buster reagent (Invitrogen), 1% (v/v) benzonase (Invitrogen), 2 mM mercaptoethanol, 10 mM EDTA, 1 mM PMSF, and an EDTA-free Complete Protease Inhibitor Cocktail tablet (Roche, Indianapolis, IN, USA). The mixture incubated at ambient temperature for 20 min on a microtiter shaker at 300 rpm (Ratek, Victoria, Australia) was centrifuged (4,500xg, 10 min, ambient temperature) to yield the first soluble fraction. The pellet was resuspended in 4% (w/v) sodium dodecyl sulphate (SDS), and after incubation (10 min, ambient temperature) the second solubilised fraction was collected (4,500xg, 10 min, ambient temperature). The pellet was solubilised in 8 M urea in 0.1 M NaOH to collect the third solubilised fraction after centrifugation (4,500xg, 10 min, ambient temperature). The fourth fraction was an insoluble material after alkali/urea treatment. Fractions 1-4 were evaluated by the immunoblot blot analyses using a mouse anti 6xHis monoclonal (Clontech, Takara Bio, Shiga, Japan) and polyclonal antibody raised against HvNIP2;1 (using the SVAADDDELDHIPV C-terminal peptide) (6), provided by the courtesy of Professor Jian Feng Ma (Okayama University, Japan). Samples containing high levels of HvNIP2;1 were analysed by SDS-PAGE and immunoblot (IB) analyses.

**Enzymatic digestion and disruption of *Pichia* cells to isolate microsomal membrane fraction.** *Pichia* cells with the highest expression of HvNIP2;1, based on the fractionation technique described above (5) were selected. Cells grown as described above were thawed on ice and digested with Zymolyase from *Arthrobacter luteus* (MPI Biomedicals, Australia) at 1 U/g of cells (7) in the 10 mM Tris-HCl buffer (pH 6.5) under rotary shaking for 2 hours at ambient temperature. Digested cells were centrifuged (500xg, 10 min, 4 ºC), and the pellet was re-suspended in a breaking buffer (BB) [50 mM sodium phosphate buffer (pH 7.5), 5% (v/v) glycerol, 1 mM DTT, 1 mM PMSF and 1 mM EDTA] and disrupted in a BeadBeater Disrupter (Biospec Products, Bartlesville, OK, USA) with acid-washed glass beads of 0.1 mm diameter (Biospec Products) using the protocol as described (5). Briefly, cells were processed *via* a ten-times disruption protocol (each lasting for 1 min), with intermittent cooling intervals (3 min), disintegrated cells centrifuged (12,000xg, 10 min, 4 ºC) and the cell debris and glass beads were suspended in 1 mL of BB, vortexed and centrifuged (12,000xg, 10 min, 4 ºC). The supernatant was transferred to polyallomer test tubes (Beckman Coulter, CA, USA) and subjected to high-speed centrifugation (200,000xg, 60 min, 4 ºC). The pellet containing the microsomal membrane fraction was resuspended in 100 µL of a storage buffer [50 mM sodium phosphate buffer (pH 7.5) 10% (v/v) glycerol, 1 mM DTT, 1 mM PMSF, 0.1 mM EDTA, 100 mM KCl, and an EDTA-free protease inhibitor cocktail (Roche)] and evaluated for the presence of HvNIP2;1 by SDS-PAGE combined with IB analyses. Alternatively, the membrane pellet was processed as described in the section below.

**Urea/alkali treatment of microsomal membrane fractions.** The membrane pellet collected by ultra-centrifugation from the last step described in the above section was treated as follows (8, 9): the pellet was resuspended in the 5 mM Tris-HCl (pH 9.5) buffer, containing 5 mM EDTA, 5 mM EGTA and 4 M urea, and contaminating proteins adhering to membranes were removed by centrifugation (100,000g, 40 min, 4 ºC). The pellet was suspended in 20 mM NaOH (approximate pH 12), incubated for 5 min at ambient temperature, centrifuged (100,000xg, 40 min, 4 ºC) and re-suspended in the 5 mM Tris-HCl (pH 8) buffer containing 2 mM EDTA, 2 mM EGTA, 100 mM NaCl, and again cleared by centrifugation (100,000xg, 40 min, 4 ºC). The final pellet containing urea/alkali-stripped microsomal plasma membranes was suspended in 100-250 µL (depending on the amounts of used cells) in the 20 mM HEPES-NaOH (pH 7.8) solubilisation buffer (SB) containing 50 mM NaCl, 10% (v/v) glycerol, 2 mM mercaptoethanol, and stored on ice to proceed with SMA solubilisation.

**Solubilisation of HvNIP2;1 from the urea/alkali-treated microsomal membrane fractions by SMA.** This step was conducted on the urea/alkali-treated microsomal membrane fraction obtained as described above, by adding SMA (10) from the 10% (w/v) stock solution in SB to 2% (w/v) concentration. Solubilisation proceeded for 4 hours under rotary shaking, after which the solubilised protein was centrifuged (200,000xg, 60 min, 4 °C) and the supernatant fraction was used for purification of the HvNIP2;1 protein. No additional SMA was added during further purification steps.

**Purification of HvNIP2;1 *via* Immobilised Metal Affinity Chromatography (IMAC).** The SMA-solubilised preparation was incubated with 0.5-1 mL of the Complete His-Tag Purification Resin (Roche, Indianapolis, IN, USA) equilibrated in SB and incubated for 16-18 hours at ambient temperature. Resin with bound protein was packed in a disposable Bio-Rad column (Hercules, CA, USA), and bound protein was eluted with 300 mM imidazole in SB at 1 mL/min flow rate at 4 °C (11). Fractions (1 mL) were analysed by SDS-PAGE combined with IB, as described above. Positive fractions were pooled and concentrated to 200 µL on a Microcon Ultracel YM10 micro-concentrator (50 kDa exclusion limit, Millipore Billerica, USA). The final preparation was aliquoted and stored with 20% (v/v) glycerol at -80 ºC.

**Reconstitution of HvNIP2;1 in liposomes.** DMPC lipids were dissolved at 10 mg/mL in chloroform, dried on a rotary evaporator under vacuum for 30 min, and rehydrated in the liposome buffer (LB) [20 mM Tris-HCl (pH 8.0) containing 100 mM KCl]. The lipid mixture was sonicated until clear and filtered through 100 nm pores (Avanti) of a uniform size using the LiposoFast Hand Extruder (Avestin, Irvine, CA, USA). The SMA-solubilised HvNIP2;1 protein preparation and filtered DMPC liposomes were mixed at a ratio of 1:50 on a weight basis, mixed by gentle shaking at room temperatures for 15 min and the mixture was dialysed in LB containing 50 mM $MgCl_2$ to disrupt the SMA polymer. The 50 mM $MgCl_2$ concentration was maintained in all the buffers after this step. No protein precipitation was observed during the reconstitution of HvNIP2;1 in DMPC liposomes.

**Isolation of homogenous DMPC liposomes with reconstituted HvNIP2;1 through floatation on the iohexol gradient.** Equal volumes (50 µL) of liposomes with reconstituted HvNIP2;1 and 80% (w/v) iohexol in the 25 mM HEPES-NaOH buffer (pH 7.5) containing 100 mM NaCl and 10% (w/v) glycerol were mixed (1, 5). The mixture was transferred to an ultra-clear polyallomer test tube (Beckman Coulter), overlaid with 350 µL of 30% (w/v) iohexol in the 25 mM HEPES-NaOH buffer (pH 7.5) containing 100 mM NaCl and 10% (w/v) glycerol, and with 100 µl of the 25 mM HEPES-NaOH buffer (pH 7.5) containing 100 mM NaCl. The mixture was centrifuged (100,000xg, 4 hours, 4 ºC) in the L-80XP ultra-centrifuge using the SW55Ti swinging-bucket rotor (Beckman Coulter). After ultra-centrifugation, 60 µl fractions were sequentially collected from the top of the gradient to the bottom and examined by SDS-PAGE and IB. Selected fractions of liposomes with embedded HvNIP2;1 were pooled, centrifuged (10,000xg, 2 min, 4 °C), resuspended in LB, dialysed for 18 hours in LB at 4 °C using 10-kD cut-off Slide-ALyserMini dialysis cups (Thermo Fischer Scientific, Rockford, IL, USA) and used in stopped-flow light scattering recordings. Fractionated proteo-liposomes stored on crushed ice remained stable for up to 5 days.

**Analytical techniques**. Protein samples were mixed with the SDS-PAGE loading buffer (5) and incubated at 37 ºC for 30 min. Samples are loaded onto a gel and run at 150 V for 90 min followed by IB analyses using 0.45 µm polyvinyl difluoride transfer membrane and detection with antibodies (5). Working solutions of antibodies were prepared in 25 mM Tris-HCl buffer, pH 7.5 containing 137 mM NaCl, 3 mM KCl, and 0.05% (w/v) Tween 20. Blots were incubated with antibodies for 1-16 hours with gentle agitation at 4-8 ºC. IB blots were developed with the Novex® ECL HRP Chemi-Luminescent Substrate Reagent Kit (Invitrogen) or with the BCIP/NBT-purple liquid reagent (Sigma-Aldrich) following the manufacturer's instructions. Proteins on SDS-PAGE gels were stained with Coomassie Brilliant Blue R-250 (Sigma-Aldrich) and the semi-quantitative estimation of protein content on SDS-PAGE gels was based on known amounts of BSA (fraction V, Sigma-Aldrich).

**Heterologous expression of HvNIP2;1 in *Xenopus laevis* oocytes and oocyte swelling**. HvNIP2;1 expression in oocytes was performed as described (12, 13). Briefly, native HvNIP2;1 DNA was inserted in the Gateway-enabled pGEMHE vector (2, 12, 13) and complementary RNA (cRNA) was transcribed using the Ambion mMESSAGE mMACHINE kit (Life Technologies, Carlsbad, CA, USA). 23 ng cRNA in 46 nL of RNA-free water or an equal volume of RNA-free water were injected in oocytes, followed by incubation in ND-96 for 24–48 hours before measurements (12, 13). Permeability of HvNIP2;1 expressed in oocytes to solutes was investigated after the transfer of oocytes to the 5-fold diluted solution of ND96 (5 mM HEPES-NaOH buffer, pH 7.4 containing 96 mM NaCl, 2 mM KCl, 1.8 mM $CaCl_2$, and 1 mM $MgCl_2$) supplemented with solutes at 160 mM concentration, which equalled to 200 mOsmol/kg osmolarity. 0.5 mM $AgNO_3$ (14) was used to inhibit the permeation of HvNIP2;1 expressed in oocytes.

**3D molecular model of HvNIP2;1 and the crystal structure of SoPIP2;1.** The full-length homology model of monomeric HvNIP2;1 was generated using the protein structure prediction tool in PHYRE2 Protein Fold Recognition Server (15), which provides a 94% confidence level at an accuracy higher than 90%. The model of HvNIP2;1 was constructed based on the structural information available before the coordinates of rice HvNIP2;1 (PDB 7nl4 and 7cjs) were released (3 November 2021) (16, 17). In these structures, respective 44 and 24(7cjs) or 22(7nl4) N- and C-terminal residues remained unresolved. The coordinates of spinach (*Spinacia oleracea*) SoPIP2;1 AQPs an open state conformation (18, 19) were taken from the Protein Data Bank (PDB accession 2B5F). PROCHECK (20) was used to evaluate the stereochemical and geometrical quality of the HvNIP2;1 model, indicating that 99.6% of residues were located in the most favourable, additionally allowed and generously allowed regions, while one residue (L290) positioned close to the C-terminus, located to a disallowed region. ProSa2003 (21) Z-scores (measures of Cβ-Cβ pair interactions) were -4.4 and -6.1 for the HvNIP2;1 model and the SoPIP2;1 crystal structure (PDB 2b5F), respectively. These parameters indicated that structures had favourable conformational energies relative to their length and were placed in the allowed Z-score conformational energy regions (21). Structural images were created in the PyMOL Molecular Vizualization System V2.8.0.6 (Schrődinger LLC, Portland, OR, USA).

**Molecular dynamics (MD) simulations of HvNIP2;1 and SoPIP2;1.** Structural relaxations and refinements of the predicted HvNIP2;1 model and the crystal structure of SoPIP2;1 progressed with all-atom MD simulations for 100 nsec, using previously developed protocols (5, 22-24). Briefly, AQPs were embedded in the POPE lipid bilayer and solvated in TIP3P water molecules (25) with $Na^+$ ions (26) to neutralise the overall charges. A membrane builder (27-29) in the CHARMM-GUI web-based server (29) was used for HvNIP2;1 and SoPIP2;1 embedded in lipid bilayers. AMBER 16 (30) with FF14SB force field (31) and CHARMM-36 parameters for lipid bilayers (32) were used for all-atom conventional MD simulations of proteins, including minimisation, equilibrations, and production runs of proteins. Energy minimization was performed with 10,000 steps for solvent molecules while proteins and lipid bilayers were harmonically restrained with 200 kcal/mol, followed by gradually heating up the system to 200 K with the same restraints followed by a short equilibration MD run under NPT for 200 ps. Additional 10,000 step minimization was applied with the 25 kcal/mol restraints on the protein and lipid bilayers followed by a second short NVT run that was carried out for 200 ps with a 25 kcal/mol restraint. The harmonic restraint was gradually reduced by applying five additional minimization steps with decreasing energy from 20, 15, 20, 5, and 0 kcal/mol. After the series of minimizations, the entire system was heated to 300 K. Production runs were carried out for 100 nsec under the NPT condition with a time step of 2 fs. To calculate long-range electrostatic potentials in all directions of periodic boundary, the Particle Mesh Ewald (PME) summation method was used (33) MD simulations protocols for water permeation by HvNIP2;1 and SoPIP2;1 proceeded according to a published protocol (34).

Residues located in pores of HvNIP2;1 and SoPIP2;1, listed below, are categorised based on these properties: black, hydrophobic aliphatic, and aromatic with neutral side chains; green, hydrophilic aliphatic side chains; red, acidic side chains; blue, basic side chains; *, unique side chains.

**Residues involved in permeation pathways of HvNIP2;1 and SoPIP2;1:**

**HvNIP2;1:** E54, S57, T58, L61, V62, T65, C66, A68, S85, G88*, Val92, H106, M107, N108, V111, T129, A132, N133, G155*, T156, T174, M177, T181, S202, V203, T206, S207, A210, G211*, A212, G216 *, S217, N219, R222, T223.

**SoPIP2;1:** F51, L52, T55, V56, I85, V89, A93, G98*, I100, N101, V104, T172, L175, V176, V179, A182, P199*, I202, V206, V209, H210, T219, G220*, I221, N222.

**Phylogenomic analyses.** Viridiplantae MIP sequences with matches to the PF00230 PFAM (35) were retrieved from Phytozome 12.1 (https://phytozome.jgi.doe.gov/pz/portal.html) (36) (Dataset S1). Archaean, bacterial, fungal, and metazoan sequences were retrieved from the UniProt reference genomes using top hits from the EBI hmmsearch implementation (https://www.ebi.ac.uk/Tools/hmmer/search/hmmsearch) (37, 38). Barley and wheat sequences were curated from in-house collections and the NCBI GenBank (39). Excessively long, short, or fragmented sequences were manually removed. Two datasets were prepared, one with only Viridiplantae and a larger expanded collection of plant and archaean, bacterial, fungal, and metazoan sequences. Jalview (40) was used to identify a 90% redundancy threshold for archaean, bacterial, fungal, and metazoan sequences and to manually select cluster representatives. The AlignSeqs function from DECIPHER (38) was used to align 164 selected Viridiplantae sequences. Clipkit (41) was used to trim excessively gapped sites under the gap model (g=0.95). The expanded all domain dataset was prepared using hmmalign where residues were assigned to the MIP PF00230 HMM from the Pfam database. The flanking unassigned residues were excluded using the hmmalign trim function.

As MIP sequences were relatively short, data could not support the fit for complex, parameter-rich models to deep alignments. Thus, maximum-likelihood analyses were restricted to Viridiplantae, while the analyses of Virdiplantae, archaean, bacterial, fungal, and metazoan sequences were limited to distance methods. Substitution model selection for the Viridiplantae and all domain data was performed using ModelTest-NG (42) with LG+G4 determined as best-fit for both data under the AICc. The phylogeny was calculated using RAxML-NG v1.0.2 (43). The best-known maximum-likelihood tree was selected based on final GAMMA scores after 150 random and

150 parsimony start-tree searches. Confidence values were determined by calculating 1000 transfer bootstrap estimate replicates (44). Distance analysis was performed with FastME 2.0 (45) using the LG+G4 substitution model. Confidence values were determined by calculating 1000 bootstrap replicates.

### Supplementary Results

**Expression, purification, and reconstitution of HvNIP2;1 in liposomes.** *Pichia* transformants were screened for high-level protein expression, where HvNIP2;1 appeared predominantly in alkaline soluble or insoluble alkaline/urea fractions (data not shown). The identity of purified HvNIP2;1 was confirmed by immunoblot, SDS-PAGE, and IB analyses using anti-His (Fig. 1) and anti-HvNIP2;1 antibody. These analyses revealed that HvNIP2;1 folded predominantly into monomeric (apparent molecular mass of around 34 kDa) and dimeric (apparent molecular mass of around 70 kDa) forms, but tetrameric forms (apparent molecular mass of around 150 kDa) were also detected (Fig. 1). Electrophoretic profiles of purified HvNIP2;1 showed diffused bands suggesting that HvNIP2;1 is N-glycosylated at putative Ser26 but occupations of O-glycosylation sites were also possible. No degradation of purified HvNIP2;1 was observed after three weeks at 4 $^{o}$C indicating that it was amenable for permeation studies.

To confirm that HvNIP2;1 was incorporated in DMPC liposomes and not simply aggregated during HvNIP2;1 reconstitution, liposomes with embedded HvNIP2;1 were subjected to floatation by iohexol density gradient ultra-centrifugation. As indicated in Fig. 1B, fractions 2-4 containing proteo-liposomes with HvNIP2;1, detected with anti-His antibody, floated near the top of the gradient, and most of HvNIP2;1 incorporated in DMPC liposomes.

**Legends for Movies S1-S24**

**Movies S1-S4 (separate files).** Molecular animation of water permeation by HvNIP2;1 *via* A-B-C (movie S1), A-B-A (movie S2), C-B-A (movie S3), and C-B-C (movie S4) directions. Water molecules are shown in blue spheres. One representative water molecule is highlighted in red to assist in examining permeation.

**Movies S5-S8 (separate files).** Molecular animation of water permeation by SoPIP2;1 *via* C-B-A (movie S5), C-B-C (movie S6), A-B-C (movie S7), and A-B-A (movie S8) directions. Water molecules are shown in blue spheres. One representative water molecule is highlighted in red to assist in examining permeation.

**Movie S9 (separate file).** Molecular animation of BA permeation by HvNIP2;1 *via* A-B-C direction. BA is shown in cpk spheres.

**Movie S10 (separate file).** Molecular animation of BA permeation by HvNIP2;1 *via* A-B-C direction with associated water molecules. BA and water molecules are shown in cpk spheres.

**Movies S11-S12 (separate files).** Molecular animation of BA permeation with associated water molecules by HvNIP2;1 (movie S11) and SoPIP2;1 (movie S12) *via* A-B-C directions. BA is shown in yellow spheres and associated water molecules are in cpk spheres, respectively.

**Movies S13-S14 (separate files).** Molecular animation of conformational changes of HvNIP2;1 (movie S13) and SoPIP2;1 (movie S14) during sucrose transport *via* C-B-A directions. Sucrose is shown in cyan cpk sticks.

**Movies S15-S16 (separate files).** Molecular animation of sucrose transport by HvNIP2;1 *via* A-B-C (movie S15) and C-B-A (movie S16) directions. Sucrose is shown in cyan cpk sticks.

**Movies S17-S20 (separate files).** Molecular animation of KCl permeation by HvNIP2;1 *via* A-B-C (movie S17) and C-B-A (movie S18) directions, and by SoPIP2; *via* A-B-C (movie S19) and C-B-A (movie S20) directions. $K^+$ and $Cl^-$ ions are shown in brown and cyan spheres, respectively.

**Movies S21-S24 (separate files).** Molecular animation of $NaNO_3$ permeation by HvNIP2;1 *via* A-B-C (movie S21) and C-B-A (movie S22) directions, and by SoPIP2; *via* A-B-C (movie S23) and C-B-A (movie S24) directions. $Na^+$ and $NO_3^-$ ions and associated water molecules (movie S21) are shown in yellow and cpk spheres, respectively.

**Legend for Dataset S1**

**Dataset S1 (separate file).** List of 3,157 Viridiplantae, archaean, bacterial, fungal, and metazoan sequences investigated in this work.

**Table S1.** Rate constants and permeability coefficients of HvNIP2;1 embedded in liposomes [a]

| **Permeant** [b] | **Structure** | **Rate constant ($s^{-1}$)** [c] | ***P* coefficient ($cm\ s^{-1}$)** |
|---|---|---|---|
| **Solute** | | | |
| Water [d] | 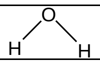 | 73.15±3.04 | $3.98\times10^{-2}\pm16.0$ |
| Boric acid | 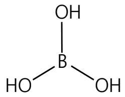 | 1.75±0.02 | $2.50\times10^{-6}\pm0.03$ |
| Germanic acid | 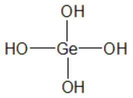 | 0.59±0.02 | $0.94\times10^{-6}\pm0.03$ |
| Urea | 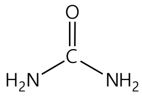 | 0.95±0.01 | $1.49\times10^{-6}\pm0.02$ |
| Glycerol | 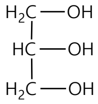 | 0.88±0.01 | $1.44\times10^{-6}\pm0.02$ |
| D-Mannitol | 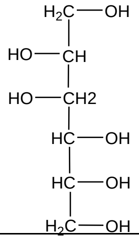 | 0.77±0.02 | $1.25\times10^{-6}\pm0.03$ |
| **Mono- and disaccharides** | | | |
| Sucrose | 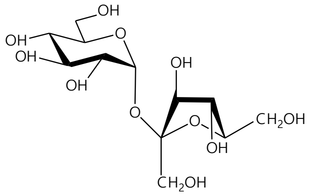 | 1.31±0.01 | $1.58\times10^{-6}\pm0.02$ |
| L-Arabinofuranose | 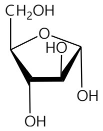 | 1.12±0.01 | $1.50\times10^{-6}\pm0.01$ |
| Lactose | 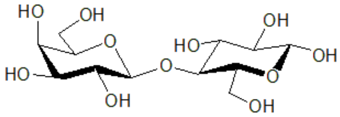 | 0.70±0.08 | $0.73\times10^{-6}\pm0.13$ |
| **Ions** | | | |
| $MgSO_4$ | 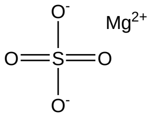 | 1.75±0.02 | $2.66\times10^{-6}\pm0.03$ |
| KCl | 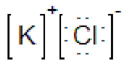 | 1.33±0.02 | $1.54\times10^{-6}\pm0.03$ |
| $CH_3COONa$ | 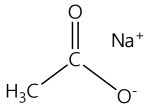 | 0.79±0.03 | $1.06\times10^{-6}\pm0.06$ |
| $NaNO_3$ | 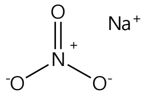 | 0.46±0.05 | $0.30\times10^{-6}\pm0.08$ |

[a] SMA-solubilised HvNIP2;1 was reconstituted in DMPC liposomes.

[b] Permeability coefficients of proteo-liposomes with HvNIP2;1 and control liposomes for water were calculated using $P_{water}$ = (V/A) × rate constant / ($V_w$ × $C_o$), and those for other solutes $P_{solute}$ = (V/A) × rate constant, where V/A is the volume to surface area ratio of liposomes (radius 50-nm), $V_w$ is the partial molar volume of water and $C_o$ is the external osmolarity after mixing. Values were rounded to two decimal points.

[c] Calculated with GraphPad Prism 9 based on two biological and two technical replicates of five averaged stopped-flow acquisitions.

[d] The $P_{water}$ coefficients ratio of proteo-liposomes with HvNIP2;1 ($3.98 \times 10^{-2}$ cm $s^{-1}$) *versus* empty liposomes ($1.66 \times 10^{-4}$ cm $s^{-1}$) is 240.

**Table S2.** Distribution of peptide motifs in bipartite segments of HvNIP2;1

| **Sequence position** | **P-value** [a] | **1st Repeat** | **Structural element** | **Sequence position** | **P-value** [a] | **2nd Repeat** | **Structural element** | **Colour in image** |
|---|---|---|---|---|---|---|---|---|
| 31-37 | 2.5e-09 | MVYYTER | N-terminal loop | 252-258 | 7.8e-09 | WTYTYIR | α-helix 6 | Yellow |
| 38-43 | 4.9e-07 | SIADYF | N-terminal loop | 116-121 | 8.8e-08 | AIFRHF | H3-HB loop | Forest green |
| 44-51 | 1.1e-08 | PPHLLKK | α-helix 1 | 263-269 | 5.4e-10 | PKDAPQK | C-terminal loop | Cyan |
| 54-66 | 2.1e-14 | EVVSTFLL VFVTC | α-helix 1 | 170-182 | 2.9e-15 | EVVVTFNM MFVTL | α-helix 4 | Magenta |
| 74-79 | 5.4e-08 | HDVTRI | Loop A | 149-154 | 2.2e-07 | HPITVI | Loop C | Blue |
| 139-144 | 9.2e-07 | CASFVL | α-helix 3 | 204-209 | 1.9e-07 | CITSIF | α-helix 5 | Red |
| 95-103 | 4.7e-08 | MIYAVGH | α-helix 2 | 187-193 | 7.0e-09 | DTRAVGE | Loop D | Black |
| 107-112 | 4.6e-08 | M**NPA**VT [b] | Re-entrant α-helix HB | 218-223 | 1.0e-08 | M**NPA****R** [b] ***T*** [c] | Re-entrant α-helix HE | Pale green/ orange |
| 123-130 | 4.9e-07 | WIQVPFY W | α-helix 3 | 230-237 | 8.8e-08 | SNRYPGLW | α-helix 6 | Grey |

[a] P-value is a measure of the false discovery rate of each analysed match (46).

[b] NPA and R222 selectivity filter residues are in underlined bold or bold, respectively.

[c] Froger's P2 T223 position is in italics bold. P1 (L148) and P3-P5 (A227, Y239, F240) are excluded.

**Table S3.** Water permeation parameters in HvNIP2;1 and SoPIP2;1 during MD simulations

| | All-through mode [a] | | | U-turn mode [a] | | Sum of water molecules (all-through mode) [a] | Sum of water molecules (U-turn mode) [b] |
|---|---|---|---|---|---|---|---|
| | A-B-C | C-B-A | Average residency time [b] | A-B-A | C-B-C | | |
| HvNIP2;1 | 8 | 23 | ~ 8 ns | 9 | 77 | 31 | 86 |
| SoPIP2;1 | 6 | 14 | ~ 4 ns | 24 | 8 | 20 | 32 |

[a] Trajectories for the last 20 nsec (40 – 60 nsec time interval).

[b] Average residency time is an approximate value (nsec) that allows a water molecule to permeate the pore.

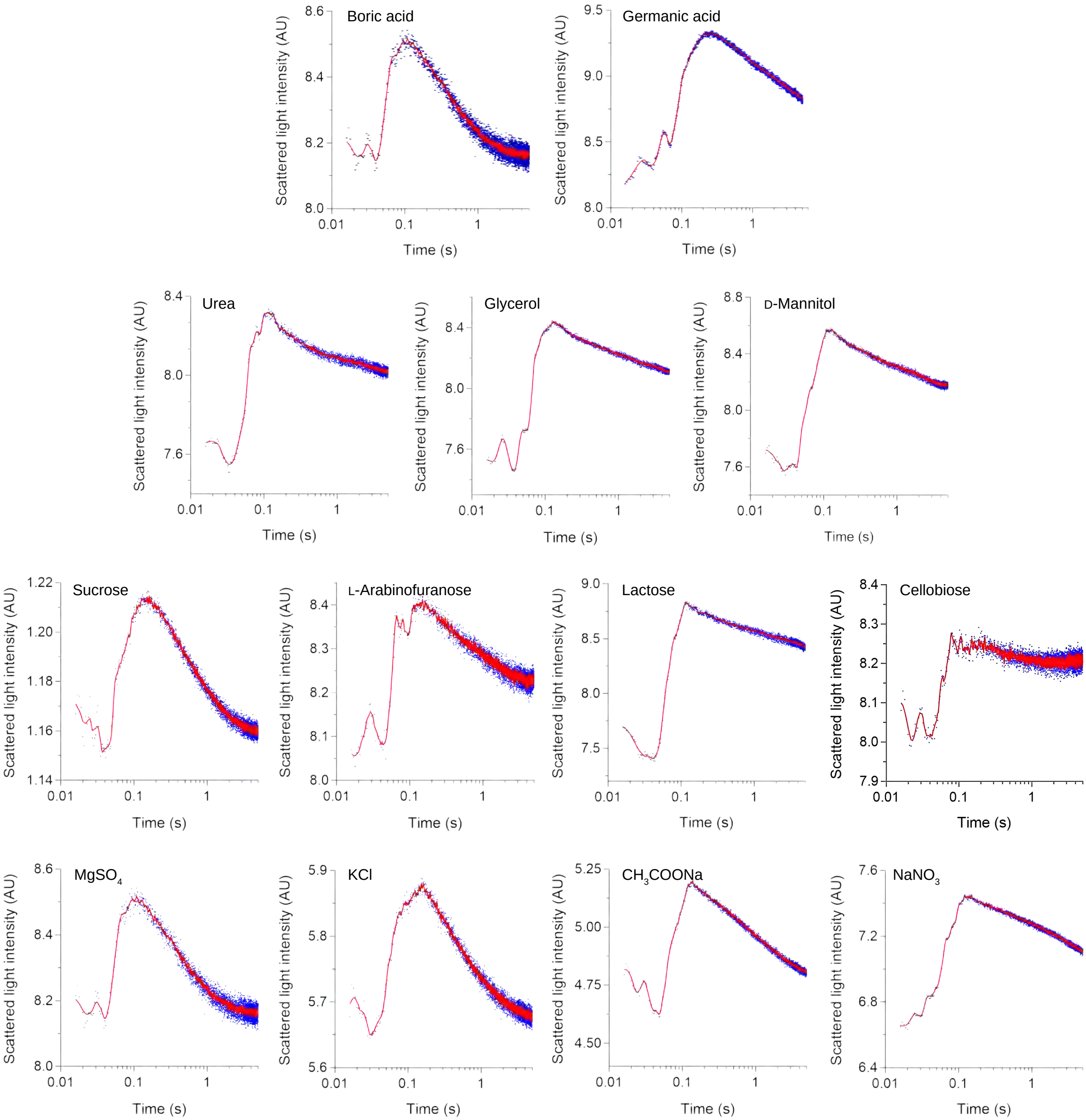
Boric acid
Germanic acid
Urea
Glycerol
D-Mannitol
Sucrose
L-Arabinofuranose
Lactose
Cellobiose
MgSO4
KCl
CH3COONa
NaNO3
Scattered light intensity (AU)
Time (s)

**Fig. S1.** Transport of permeants by HvNIP2;1 embedded in liposomes.

DMPC liposomes with embedded HvNIP2;1 were exposed to gradients of permeants generating osmotic gradients with BA and germanic acid (top panels), urea, glycerol, and D-mannitol (top-middle panels), disaccharides sucrose and lactose and a monosaccharide L-arabinofuranose (bottom-middle panels) and $MgSO_4$, KCl, $CH_3COONa$ and $NaNO_3$ (bottom panels). Uptake of permeants was measured by stopped-flow spectrophotometry, where light scattering due to the initial vesicle shrinkage (initial peak) and subsequent swelling (slope) were monitored for five seconds. Light scattering traces of five averaged traces of each permeant representing the raw (blue trace) and smoothed (red trace) data were plotted in arbitrary units (AU) in GraphPad Prism 9.

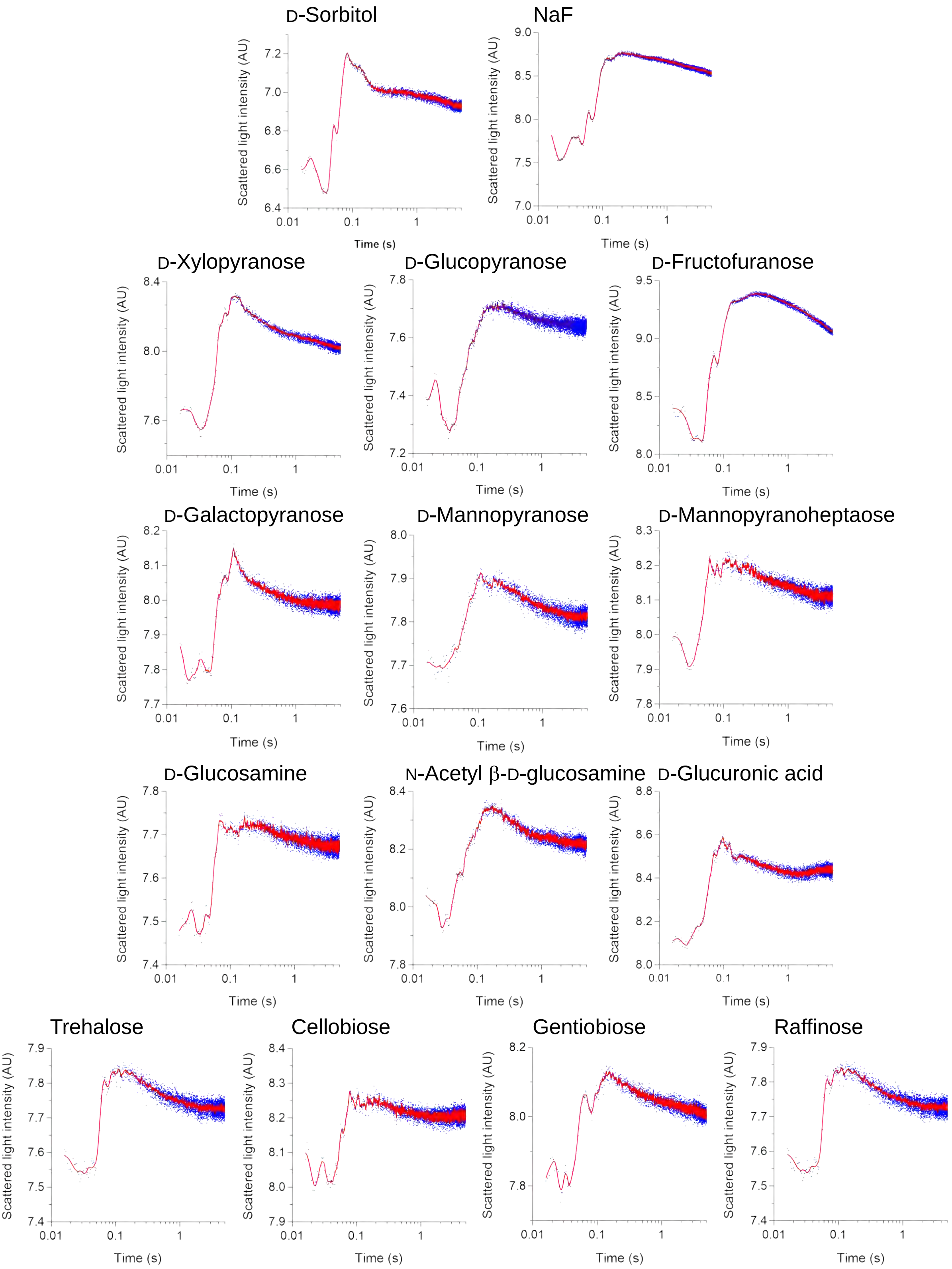

D-Sorbitol
NaF
D-Xylopyranose
D-Glucopyranose
D-Fructofuranose
D-Galactopyranose
D-Mannopyranose
D-Mannopyranoheptaose
D-Glucosamine
N-Acetyl β-D-glucosamine
D-Glucuronic acid
Trehalose
Cellobiose
Gentiobiose
Raffinose
Scattered light intensity (AU)
Time (s)
0.01
0.1
1

**Fig. S2.** Transport of permeants by HvNIP2;1 embedded in liposomes.

DMPC liposomes with embedded HvNIP2;1 were exposed to gradientso gradients of solutes generating osmotic gradients with D-sorbitol, NaF (top panels), monosaccharides D-xylopyranose, D-glucopyranose, D-fructofuranose, D-galactopyranose, D-mannopyranose and D-mannopyrano-heptaose (two top-middle panels), derivatised monosaccharides D-glucosamine, N-acetyl β-D-glucosamine, and D-glucuronic acid (bottom-middle panel), and disaccharides trehalose, cellobiose, gentiobiose, and trisaccharide raffinose (bottom panels). The uptake of permeants was measured by stopped-flow spectrophotometry, as described in Fig. 2. Light scattering traces of five averaged traces for each permeant representing raw (blue trace) and smoothed (red trace) data were plotted in arbitrary units (AU) in GraphPad Prism 9.

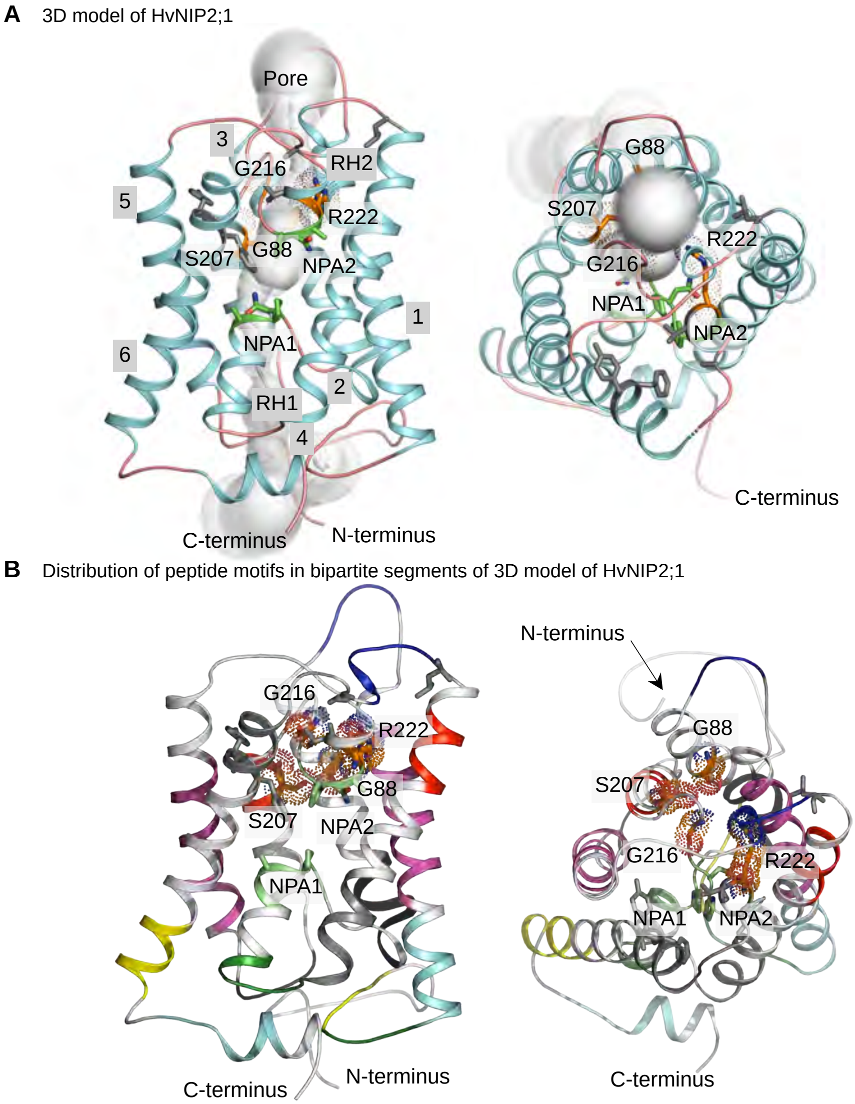


**Fig. S3.** Structural model of HvNIP2;1.

*(A)* Cartoon representation of monomeric HvNIP2;1 in two orthogonal orientations (rotated by approximately 90°) encloses a pore, shaped by the sequence of grey spheres (pore radii equal to sphere diameters). Selectivity filter residues G88, S207, G216 and R222 constituting the GSGR motif (orange sticks with dots) are positioned near the entrance to the pore, while residues of two conserved NPA1- and NPA2-motifs (green sticks) are located deeper in the pore. Residues in Froger's P1-P5 positions (L148, T223, A227, Y239, and F240) are shown in grey sticks. Numbering of α-helices 1-6 and re-entrant α-helices (RH1 and RH2) is indicated.

*(B)* The bipartite symmetry distribution of peptide motifs in two orthogonal orientations (rotated by approximately 90°). Motif pairs and their positions in secondary structural elements are coloured identically in each half of the structure. Selectivity filter residues G88, S207, G216 and R222 (orange sticks with dots), residues in NPA1- and NPA2-motifs (green sticks) and in Froger's P1-P5 positions are indicated. Six N- and 20 C-terminal residues are omitted for clarity in panels A-B.

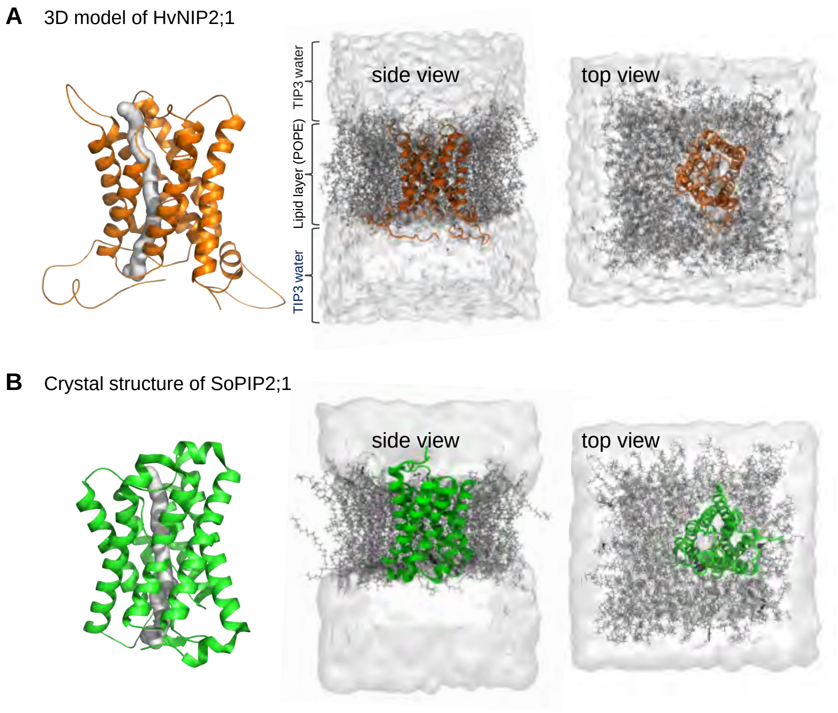


**Fig. S4.** Representative images of the structural model of HvNIP2;1 *(A)* and the crystal structure of SoPIP2;1 *(B)*. Pore morphologies are indicated in grey tubes.

Structures of AQPs embedded in POPE lipid bilayers and solvated in TIP3 water molecules are seen in two orthogonal orientations (side and top views). Atomistic models were developed in NAMD 2.12 as described in Methods.

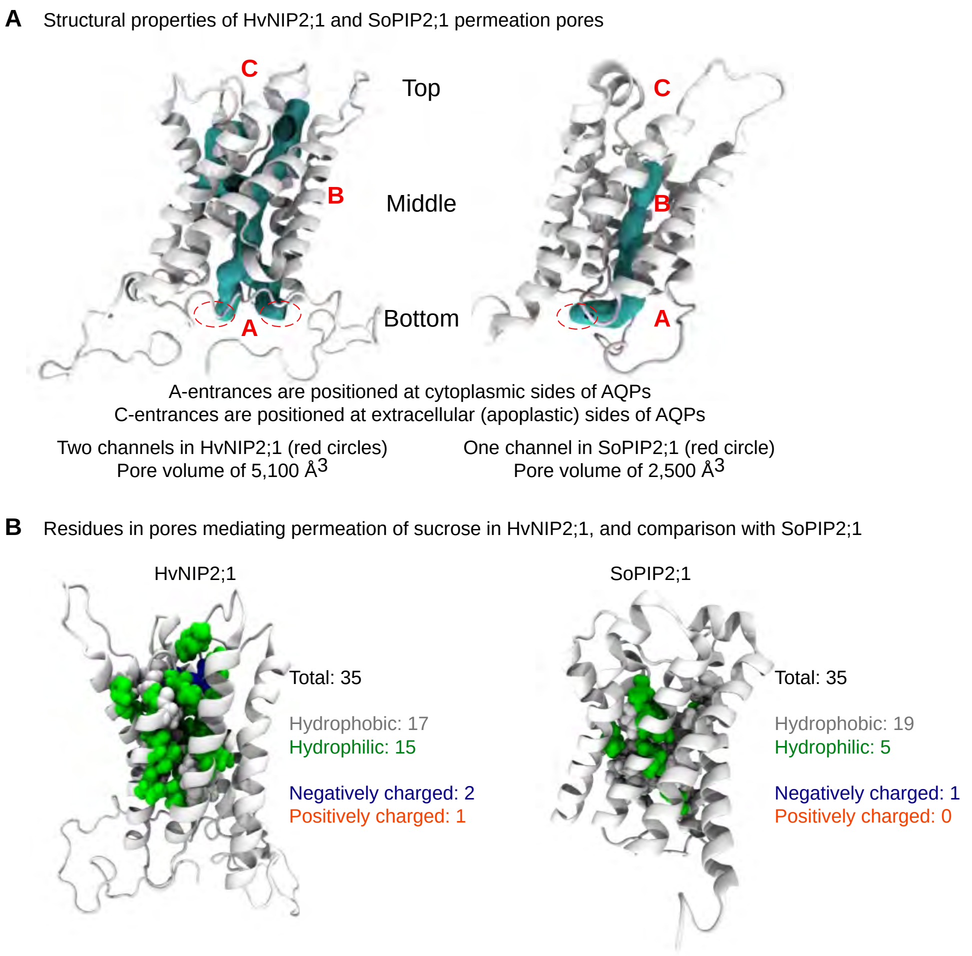


**Fig. S5.** Structural characteristics of HvNIP2;1 and SoPIP2;1 AQPs.

*(A)* Cartoon representations and morphologies of pores with delineations of three regions used in steered MD simulations: A (bottom), B (middle) and C (top). Approximate pore volumes of HvNIP2;1 and SoPIP2;1 are indicated in Å$^3$.

*(B)* Cartoon representations and properties of residues involved in sucrose permeation by HvNIP2;1 that highlight differences in pore residues allowing permeation of sucrose by HvNIP2;1 at high rates (left) while permeation of sucrose by SoPIP2;1 is unfavorable (right).

Water permeation by HvNIP2;1 and SoPIP2;1

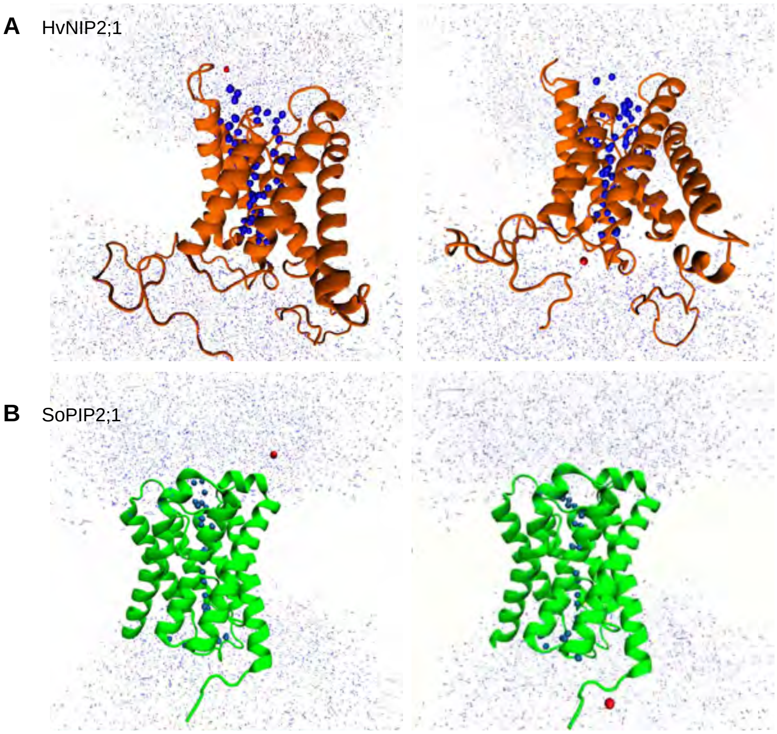


**Fig. S6.** Cartoons of HvNIP2;1 *(A)* and SoPIP2;1 *(B)* AQPs at various stages of water permeation examined *via* steered MD simulations.

These images relate to movies, in which representative water molecules (red spheres) exit at cytoplasmic or periplasmic sides of AQPs. Sizes of these representative water molecules are exaggerated for better interpretations.

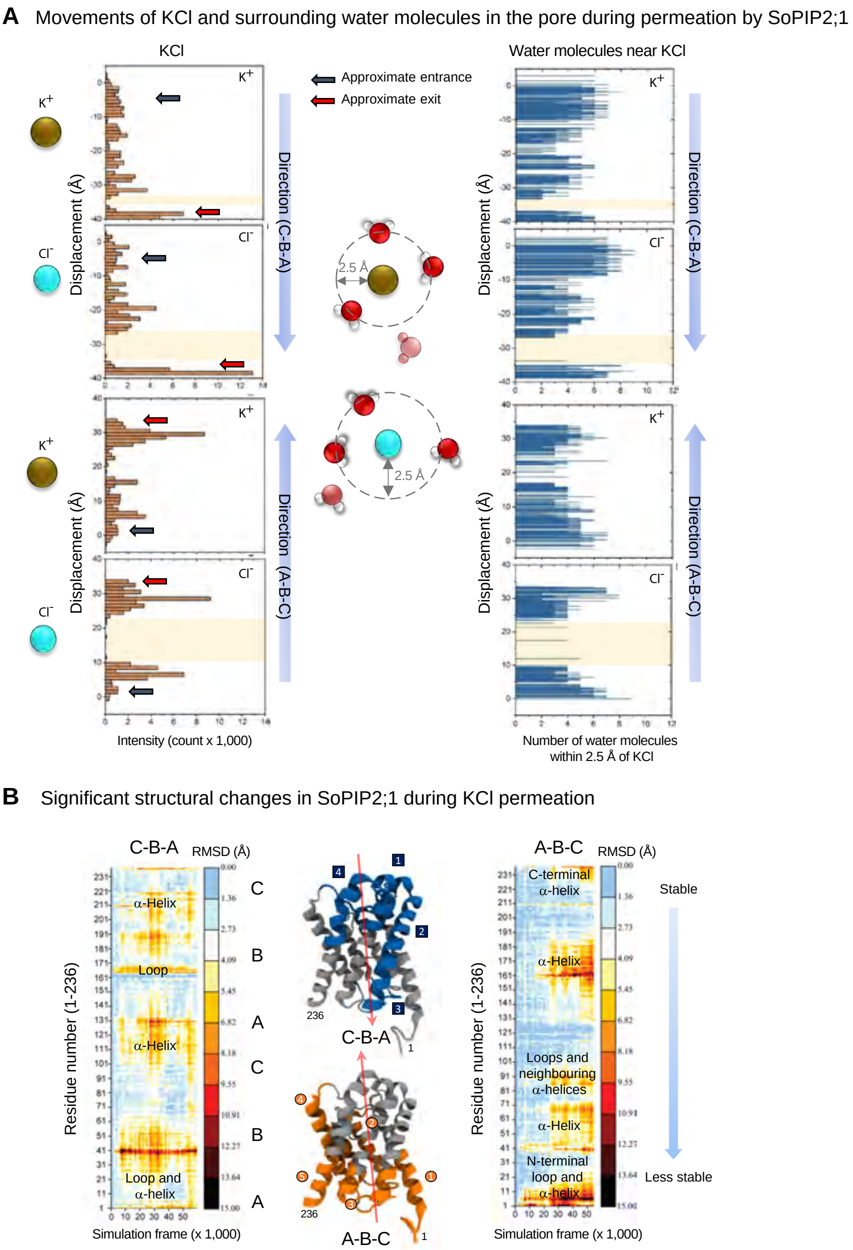
A Movements of KCl and surrounding water molecules in the pore during permeation by SoPIP2;1
KCl
Water molecules near KCl
Approximate entrance
Approximate exit
K+
Cl-
Displacement (Å)
Direction (C-B-A)
Direction (A-B-C)
2.5 Å
Intensity (count x 1,000)
Number of water molecules within 2.5 Å of KCl
B Significant structural changes in SoPIP2;1 during KCl permeation
C-B-A
A-B-C
RMSD (Å)
Residue number (1-236)
Simulation frame (x 1,000)
α-Helix
Loop
Loop and α-helix
C-terminal α-helix
Loops and neighbouring α-helices
N-terminal loop and α-helix
Stable
Less stable

**Fig. S7.** KCl permeation characteristics of SoPIP2;1 based on steered MD simulations.

*(A)* Left panels: Trace maps for $K^+$ and $Cl^-$ ions travelling through and interacting with pore residues *via* C-B-A and A-B-C directions. Intensity correlates with the likelihood of ions staying in a particular place in a pore, and displacement denotes the path of ions from starting points. Regions with sudden motions of ions through pores (not accompanied by water molecules) are highlighted in light orange. Right panels: Trace maps of water molecules (at a distance of ≤2.5 Å) surrounding $K^+$ and $Cl^-$ ions that are transported through pores. Regionslacking surrounding water molecules are highlighted and broadly match those in left panels.

*(B)* Structural changes of SoPIP2;1 during KCl permeation *via* C-B-A and A-B-C directions. Heat maps illustrate changes in residual steered MD values of superposed structures before and after permeation. Blue (C-B-A) or orange (A-B-C) colours in cartoon representations denote regions with high structural changes and those correspond to black, yellow, and red areas in heat maps, while grey colour denotes regions with low structural changes and those correspond to white and blue areas in heat maps.

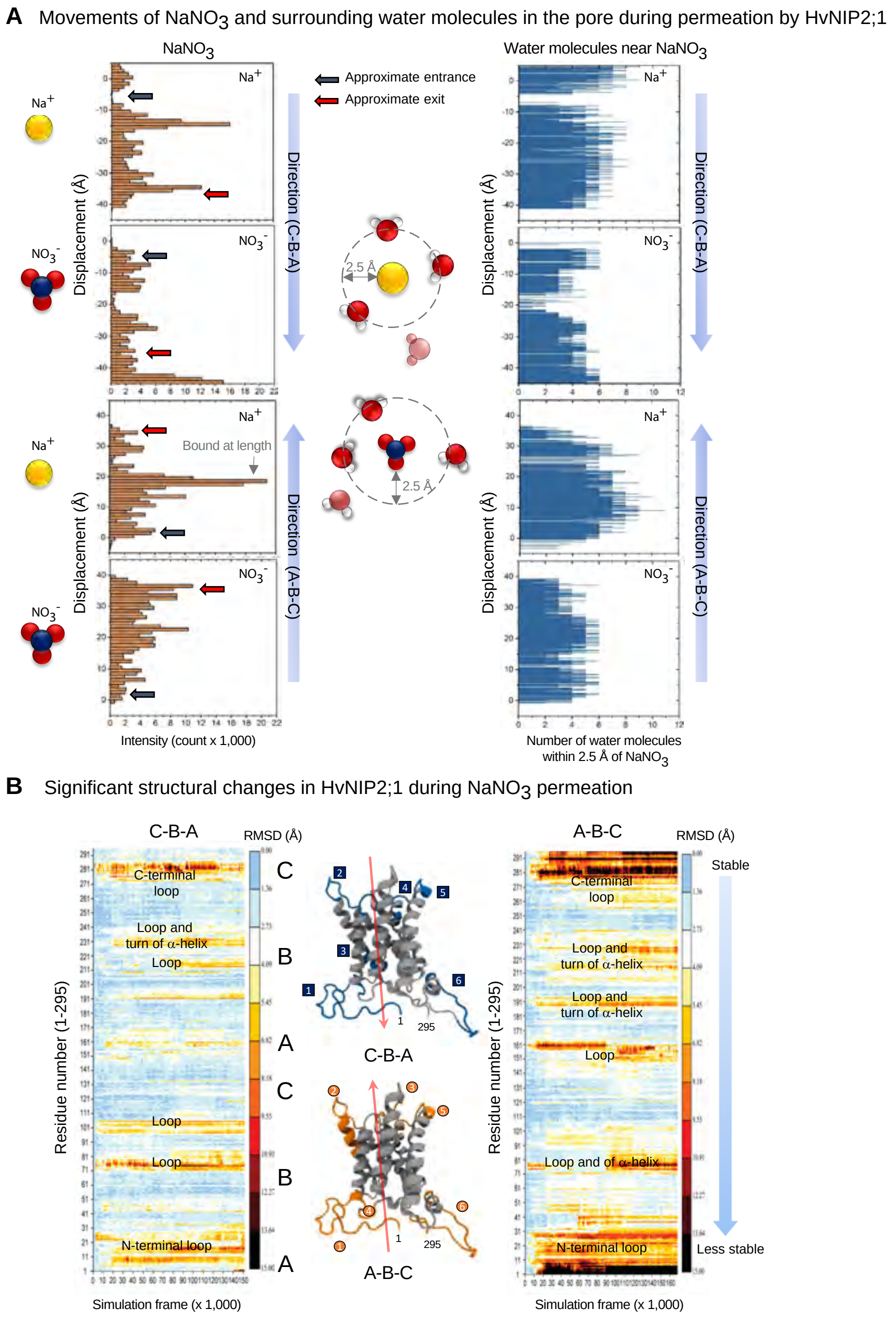
A Movements of NaNO3 and surrounding water molecules in the pore during permeation by HvNIP2;1
NaNO3
Water molecules near NaNO3
Na+
NO3-
Approximate entrance
Approximate exit
Direction (C-B-A)
Direction (A-B-C)
Displacement (Å)
2.5 Å
Bound at length
Intensity (count x 1,000)
Number of water molecules
within 2.5 Å of NaNO3
B Significant structural changes in HvNIP2;1 during NaNO3 permeation
C-B-A
A-B-C
RMSD (Å)
C-terminal
loop
Loop and
turn of α-helix
Loop
N-terminal loop
Loop and of α-helix
Residue number (1-295)
Simulation frame (x 1,000)
Stable
Less stable
C
B
A
1
295

**Fig. S8.** $NaNO_3$ permeation characteristics of HvNIP2;1 based on steered MD simulations.

(*A*) Left panels: Trace maps for $Na^+$ and $NO_3^-$ ions that travel through and interact with pore residues via C-B-A and A-B-C directions. Intensity correlates with a likelihood of ions staying in a particular place in the pore, and displacement denotes the path of ions from starting points. Regions with sudden motions of ions through pores (not accompanied by water molecules) are highlighted in light orange. Right panels: Trace maps of water molecules (at distance of ≤2.5 Å) surrounding $Na^+$ and $NO_3^-$ ions that are transported through pores. Regions lacking surrounding waters are highlighted and broadly match those in left panels.

*(B)* Significant structural changes of HvNIP2;1 during $NaNO_3$ permeation via C-B-A and A-B-C directions. Heat maps illustrate changes in residual steered MD values of superposed structures before and after permeation. Blue (C-B-A) or orange (A-B-C) colours in cartoon representations denote regions with high structural changes and those correspond to black, yellow, and red areas in heat maps, while grey colour denotes regions with low structural changes and those correspond to white and blue areas in heat maps.

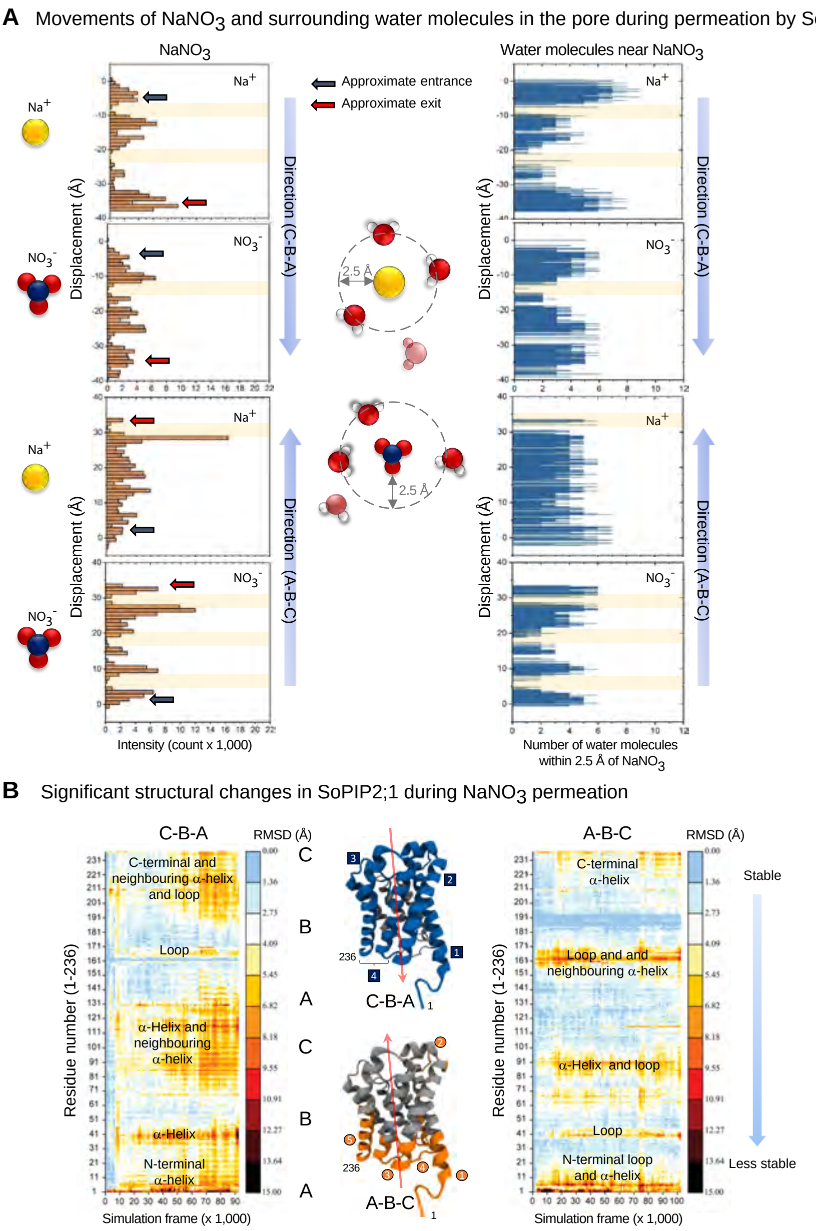
A Movements of NaNO3 and surrounding water molecules in the pore during permeation by SoPIP2;1
NaNO3
Water molecules near NaNO3
Approximate entrance
Approximate exit
Na+
NO3-
2.5 Å
Displacement (Å)
Direction (C-B-A)
Direction (A-B-C)
Intensity (count x 1,000)
Number of water molecules within 2.5 Å of NaNO3
B Significant structural changes in SoPIP2;1 during NaNO3 permeation
C-B-A
A-B-C
RMSD (Å)
C-terminal and neighbouring α-helix and loop
Loop
α-Helix and neighbouring α-helix
α-Helix
N-terminal α-helix
C-terminal α-helix
Loop and and neighbouring α-helix
α-Helix and loop
N-terminal loop and α-helix
Residue number (1-236)
Simulation frame (x 1,000)
Stable
Less stable
236

**Fig. S9.** $NaNO_3$ permeation characteristics of SoPIP2;1 based on steered MD simulations.

*(A)* Left panels: Trace maps for $Na^+$ and $NO_3^-$ ions travelling through and interacting with pore residues *via* C-B-A and A-B-C directions. Intensity correlates with the likelihood of ions staying in a particular place in the pore, and displacement denotes the path of ions from starting points. Regions with the sudden motions of ions through pores, not accompanied by water molecules, are highlighted in light orange. Right panels: Trace maps of water molecules (at a distance of ≤2.5 Å) surrounding $Na^+$ and $NO_3^-$ ions that are transported through pores. Regions lacking surroundi-ng water molecules are highlighted and broadly match those in left panels.

*(B)* Structural changes of SoPIP2;1 during $NaNO_3$ permeation *via* C-B-A and A-B-C directions. Heat maps illustrate changes in residual steered MD values of superposed structures before and after permeation. Blue (C-B-A) or orange (A-B-C) colours in cartoon representations denote regions with high structural changes and those correspond to black, yellow, and red areas in heat maps, while grey colour denotes regions with low structural changes and those corres-pond to white and blue areas in heat maps.

**A** HvNIP2;1 permeating water molecules in C-B-A direction in eight stages (bottom-middle-top)

Water molecules in the pore near NPA motifs of HvNIP2;1 and interacting residues

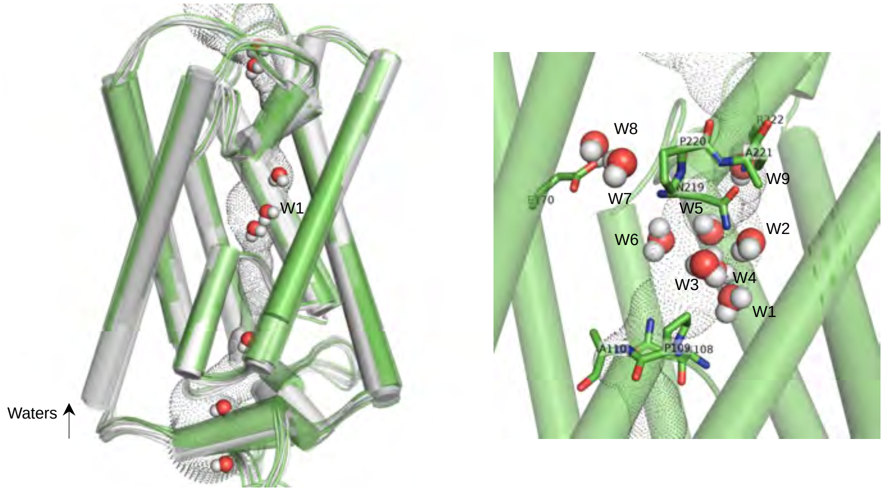


**B** HvNIP2;1 permeating BA in A-B-C direction in three stages (bottom-middle-top)

BA interacting with water molecules in the pore of HvNIP2;1 (middle position)

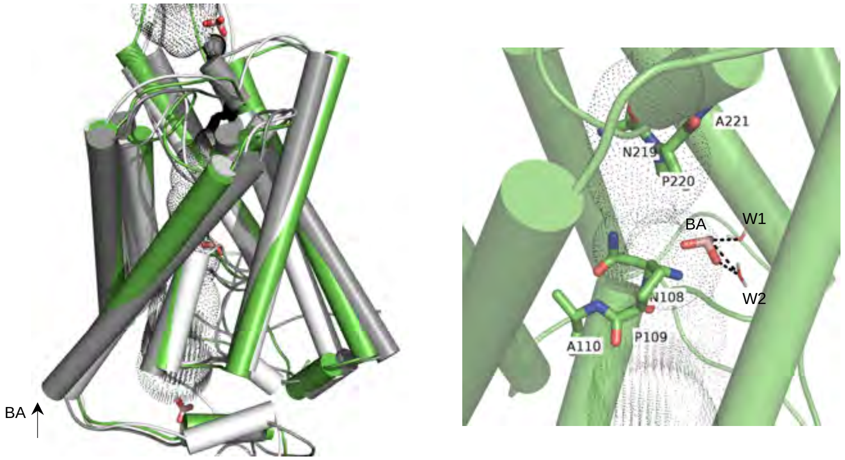


**C** HvNIP2;1 permeating sucrose in C-B-A direction in three stages (top-middle-bottom)

Sucrose interacting with water molecules in the pore of HvNIP2;1 (middle position)

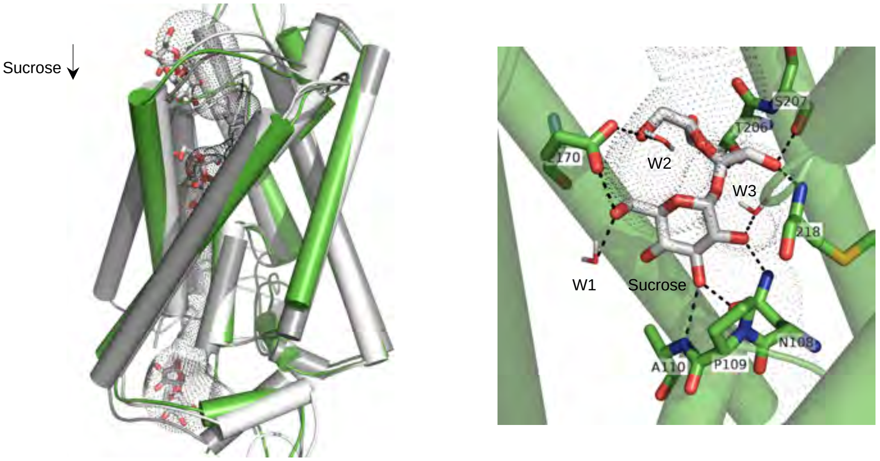

**D** HvNIP2;1 permeating KCl in A-B-C direction in three stages (bottom-middle-top)

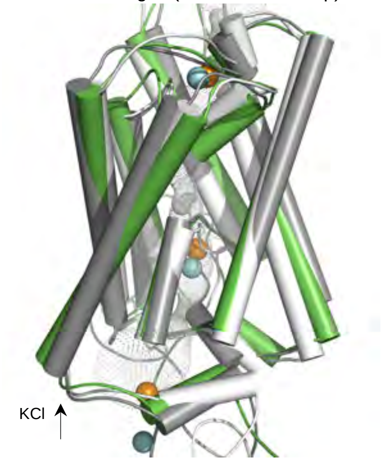


KCl interacting with water molecules in the pore of HvNIP2;1 (middle position)

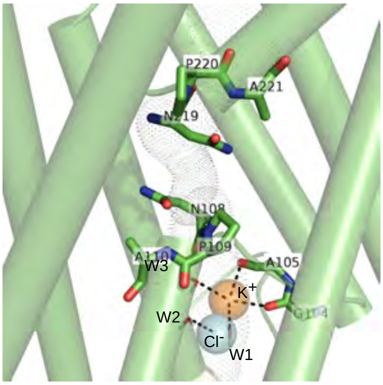


**E** HvNIP2;1 permeating KCl in C-B-A direction in three stages (top-middle-bottom)

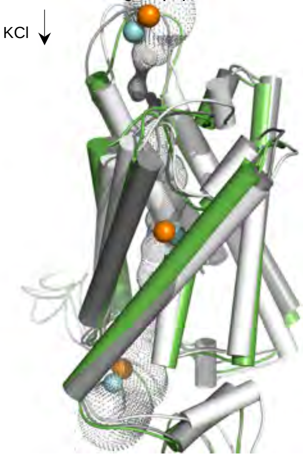


KCl interacting with water molecules in the pore of HvNIP2;1 (middle position)

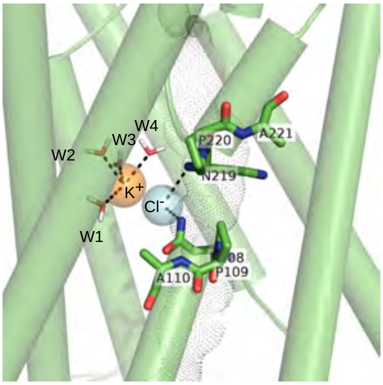


**F** HvNIP2;1 permeating $NaNO_3$ in C-B-A direction in three stages (top-middle-bottom)

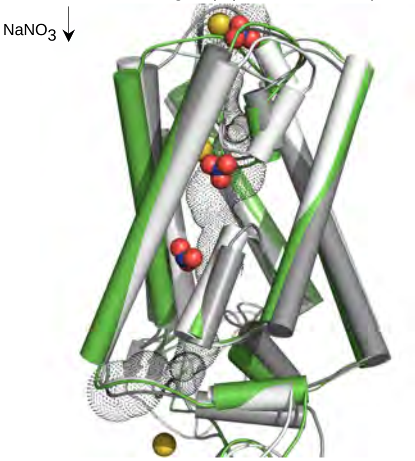


$NaNO_3$ interacting with water molecules in the pore of HvNIP2;1 (middle position)

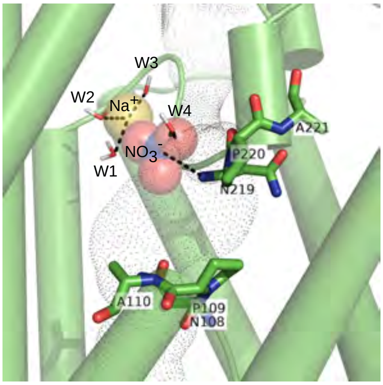

**G** SoPIP2;1 permeating water moleculess in C-B-A direction in six stages (bottom-middle-top)

Water molecules in the pore near NPA motifs of SoPIP2;1 and interacting residues

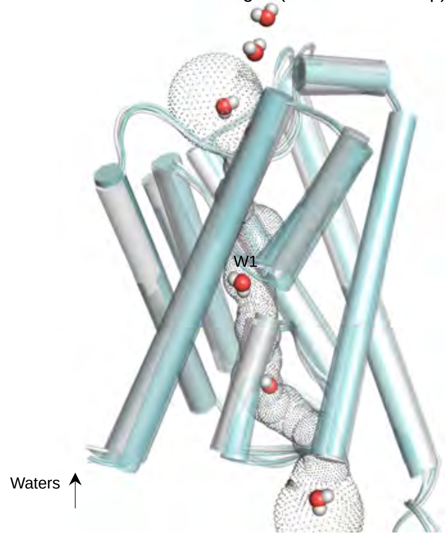


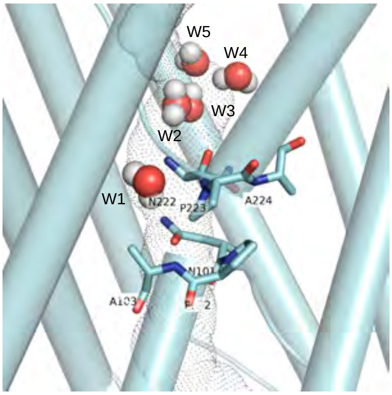


**H** SoPIP2;1 with sucrose in A-B-C direction in three stage (bottom-middle-top), where sucrose cannot be permeated

Sucrose binding water molecules in the pore of SoPIP2;1 but can not permeate the pore in A-B-C direction

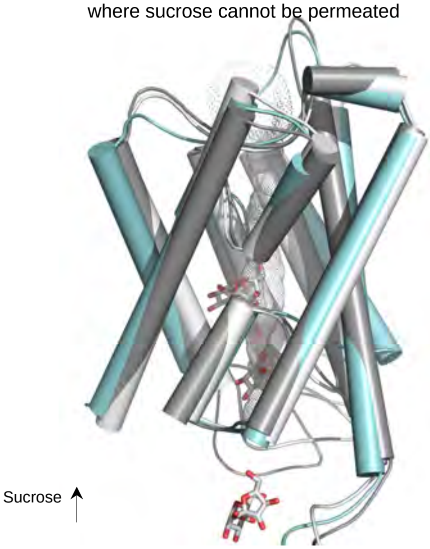


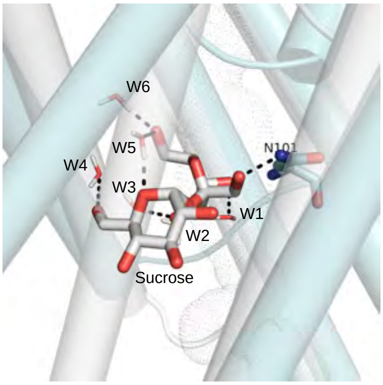


**I** SoPIP2;1 with sucrose in C-B-A direction in three stage (top-middle-bottom), where sucrose cannot be permeated

Sucrose binding water molecules in the pore of SoPIP2;1 but can not permeate the pore in C-B-A direction

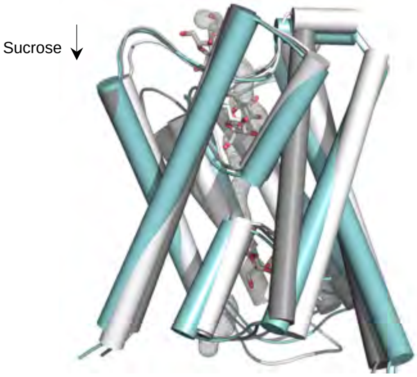


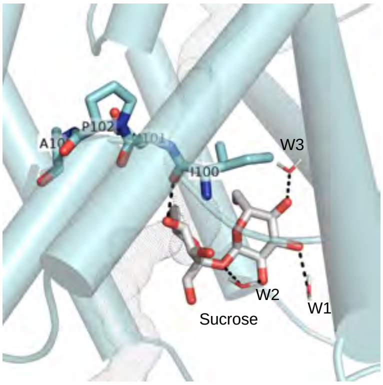

**J**

SoPIP2;1 permeating KCl in A-B-C direction
in three stages (bottom-middle-top)

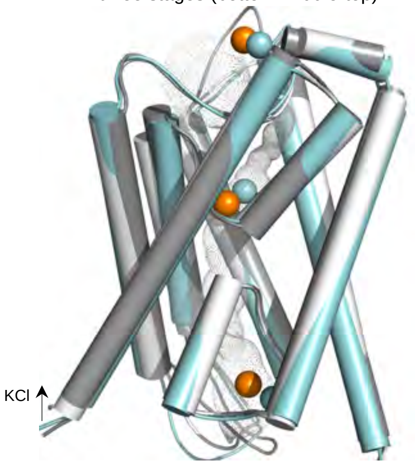


KCl interacting with water molecules
in the pore of SoPIP2;1 (middle position)

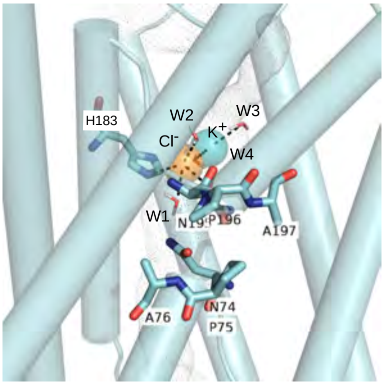


**K**

SoPIP2;1 permeating KCl in C-B-A direction
in three stages (top-middle-bottom)

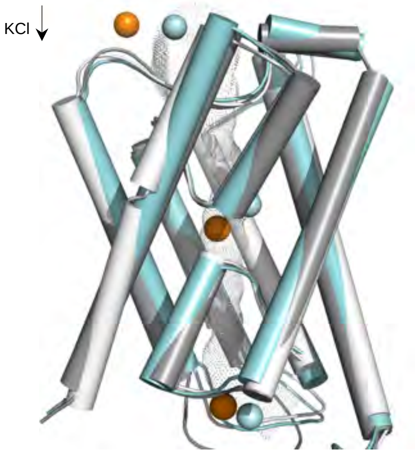


KCl interacting with water molecules
in the pore of SoPIP2;1 (middle position)

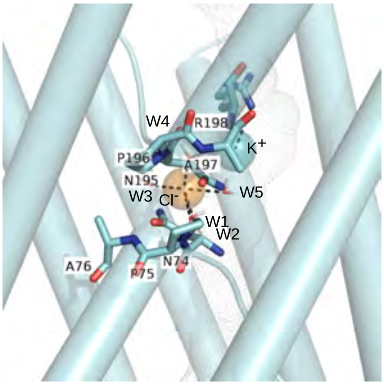


**L**

SoPIP2;1 permeating $NaNO_3$ in A-B-C direction
in three stages (top-middle-bottom)

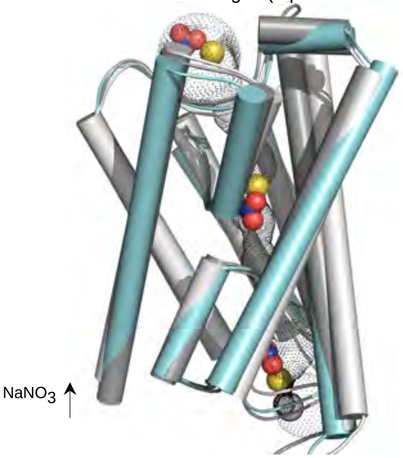


$NaNO_3$ interacting with water molecules
in the pore of SoPIP2;1 (middle position)

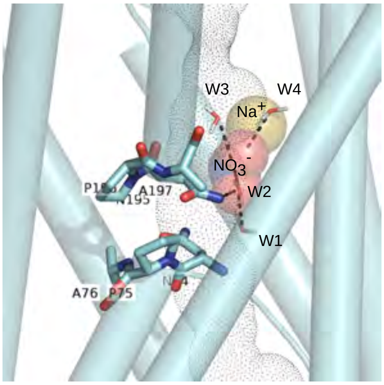

**Fig. S10.** Representative images of structures of HvNIP2;1 and SoPIP2;1 permeating water, BA, sucrose, and KCl and $NaNO_3$, developed through steered MD.

*(A)* **Left panel:** HvNIP2;1 permeating water (spheres in cpk colours) in C-B-A direction (denoted by an arrow) showing eight snapshots in various permeation stages that are superposed with RMSD values between 0.8-1.3 Å. Geometry of pores depicted in dots in left and right panels were calculated as described in Methods.
**Right panel:** Water molecule W1 (marked in the left panel) with eight neighbouring water molecules is positioned in the pore. NPA1- and NPA2-motif residues on re-entrant α-helices and Glu170 andArg222 contacting water molecules are also shown.

*(B)* **Left panel:** HvNIP2;1 permeating BA (sticks in cpk colours) in A-B-C direction (denoted by an arrow) in three stages: bottom (white), middle (green) and top (grey); three structures are superposed with RMSD values between 2.3 and 2.8 Å.
**Right panel:** BA with associated water molecules (cpk and green cpk sticks) are shown in the pore of HvNIP2;1. NPA1- and NPA2-motif residues on re-entrant α-helices are also shown. Black dashed lines denote separations of 2.7 to 3.1 Å between BA and water molecules.

*(C)* **Left panel:** HvNIP2;1 permeating sucrose (sticks in cpk colours) in C-B-A direction (denoted by an arrow) in three stages: top (white), middle (green) and bottom (grey); three structures are superposed with RMSD values between 1.9 and 2.3 Å.
**Right panel:** Sucrose with associated water molecules and residues (cpk and green cpk sticks) is shown in the pore of HvNIP2;1 in the middle position. NPA1- and NPA2-motif residues on re-entrant α-helices are also shown. Black dashed lines denote separations of 2.7 to 3.1 Å between residues, sucrose, and water molecules.

*(D)* **Left panel:** HvNIP2;1 permeating KCl (spheres in cpk colours) in A-B-C direction (denoted by an arrow) in three stages: bottom (white), middle (green) and top (grey); three structures are superposed with RMSD values between 2.0 to 2.4 Å.
**Right panel:** KCl with associated water molecules and residues (cpk spheres and sticks, and green cpk sticks) are shown in the pore of HvNIP2;1 in the middle position. NPA1- and NPA2-motif residues on re-entrant α-helices are also shown. Black dashed lines denote separations of 2.7 to 3.0 Å between residues, KCl, and water molecules.

*(E)* **Left panel:** HvNIP2;1 permeating KCl (sticks in cpk colours) in C-B-A direction (denoted by an arrow)in three stages: top (white), middle (green) and bottom (grey); three structures are superposed with RMSD values between 2.0 and 2.4 Å.
**Right panel:** KCl with associated water molecules (cpk spheres and sticks, and green cpk sticks) are shown in the pore of HvNIP2;1 in the middle position. NPA1- and NPA2-motif residues on re-entrant α-helices are also shown. Black dashed lines denote separations of 2.6 to 3.2 Å between residues, KCl, and water molecules.

*(F)* **Left panel:** HvNIP2;1 permeating $NaNO_3$ (sticks in cpk colours) in C-B-A direction (denoted by an arrow) in three stages: top (white), middle (green) and bottom (grey); three structures are superposed with RMSD values between 1.7 and 2.4 Å.
**Right panel:** $NaNO_3$ with associated water molecules (cpk spheres and sticks, and green cpk sticks) are shown in the pore of HvNIP2;1 in the middle position. NPA1- and NPA2-motif residues on re-entrant α-helices are also shown. Black dashed lines denote separations of 2.2 to 3.0 Å between residues, $NaNO_3$, and water molecules.

**Fig. S10. (cont.)** Representative images of structures of HvNIP2;1 and SoPIP2;1 permeating water, BA, sucrose, and KCl and $NaNO_3$, developed through steered MD.

*(G)* **Left panel:** SoPIP2;1 permeating water (spheres in cpk colours) in C-B-A direction (denoted by an arrow) in seven stages; seven structures in various permeation stages are superposed with RMSD values between 0.7-1.1 Å.

**Right panel:** Water molecule 1 (marked in the left panel) with four neighbouring water molecules positioned in the pore. NPA1- and NPA2-motif residues on re-entrant α-helices are also shown.

*(H)* **Left panel:** SoPIP2;1 permeating sucrose (sticks in cpk colours) in A-B-C direction (denoted by an arrow) in three stages: bottom (white), middle (cyan) and top (grey); three structures are super-posed with RMSD values between 1.7 and 2.8 Å.

**Right panel:** Sucrose with associated water molecules and residues (cpk and cyan cpk sticks) immobilised in the pore of SoPIP2;1 are shown in the middle position. Black dashed lines denote separations of 2.6 to 2.9 Å between residues, sucrose, and water molecules.

*(I)* **Left panel:** SoPIP2;1 permeating sucrose (sticks in cpk colours) in C-B-A direction (denoted by an arrow)in three stages: top (white), middle (cyan) and bottom (grey); three structures are superposed with RMSD values between 2.2 and 3.1 Å.

**Right panel:** Sucrose with associated water molecules (cpk and cyan cpk sticks) are shown in the pore of SoPIP2;1 in the middle position. NPA1-motif residues on the re-entrant α-helix are also shown. Black dashed lines denote separations of 2.6 to 2.9 Å between residues, sucrose, and water molecules.

*(J)* **Left pane**l: SoPIP2;1 permeating KCl (spheres in cpk colours) in A-B-C direction (denoted by an arrow) in three stages: bottom (white), middle (cyan) and top (grey); three structures are super-posed with RMSD values between 0.9 and 1.0 Å.

**Right panel:** KCl with associated water molecules (cpk spheres and sticks, and cyan cpk sticks) are shown in the pore of SoPIP2;1 in the middle position. NPA1- and NPA2-motif residues on re-entrant α-helices are also shown. Black dashed lines denote separations of 2.7 to 3.3 Å between residues, KCl, and water molecules.

*(K)* **Left panel:** SoPIP2;1 permeating KCl (spheres in cpk colours) in C-B-A direction (denoted by an arrow)in three stages: top (white), middle (cyan) and bottom (grey); three structures are super-posed with RMSD values between 1.2 and 1.4 Å.

**Right panel:** KCl with associated water molecules and residues (cpk spheres and sticks, and cyan cpk sticks) are shown in the pore of SoPIP2;1 in the middle position. NPA1- and NPA2-motif residues on re-entrant α-helices are also shown. Black dashed lines denote separations of 2.6 to 2.9 Å between residues, KCl, and water molecules.

*(L)* **Left panel:** SoPIP2;1 permeating $NaNO_3$ (spheres in cpk colours) during A-B-C direction (denoted by an arrow)in bottom (white), middle (cyan) and top (grey); three structures are super-posed with RMSD values between 1.5 and 1.6 Å.

**Right panel:** $NaNO_3$ with associated water molecules and residues (cpk spheres and sticks, and cyan cpk sticks) are shown in the pore of SoPIP2;1 in the middle position. NPA1- and NPA2-motif residues on re-entrant α-helices are also shown. Black dashed lines denote separations of 2.9 to 3.1 Å between residues, $NaNO_3$, and water molecules.

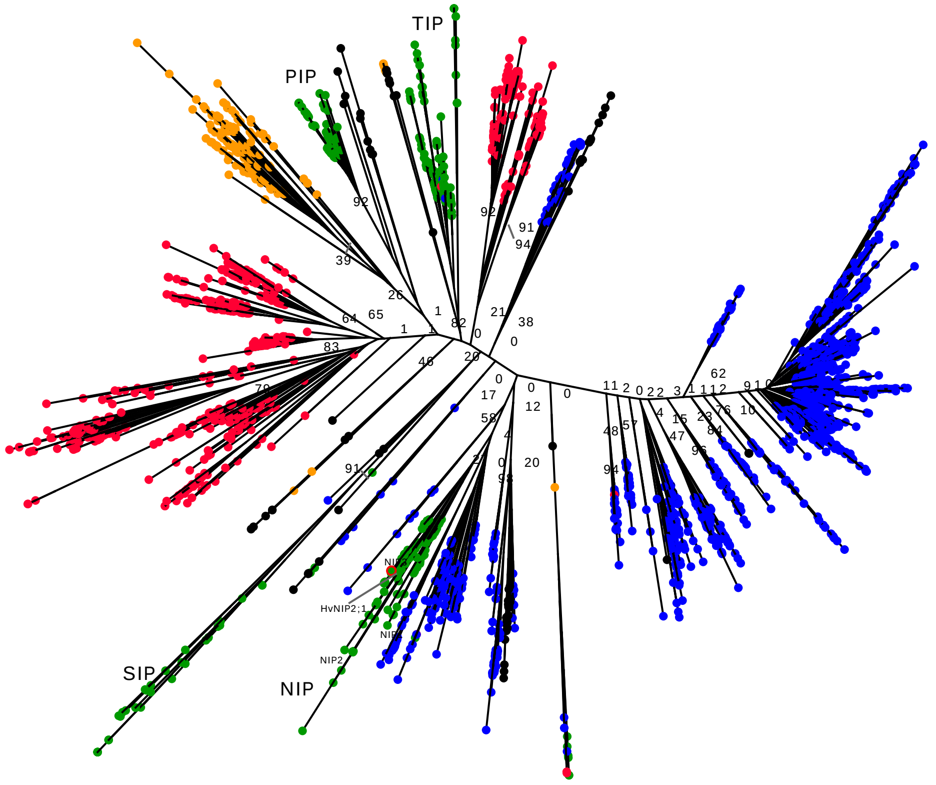


**Fig. S11.** FastME distance tree of 3,157 PF00230 MIP proteins.

Terminal nodes are colour-coded by kingdom: green, Viridiplantae; black, Archaea; blue, Bacteria; yellow, Fungi; red, Metazoa. The list of individual 3,157 entries is specified in Dataset S1. Bootstrap support values are indicated at major nodes.

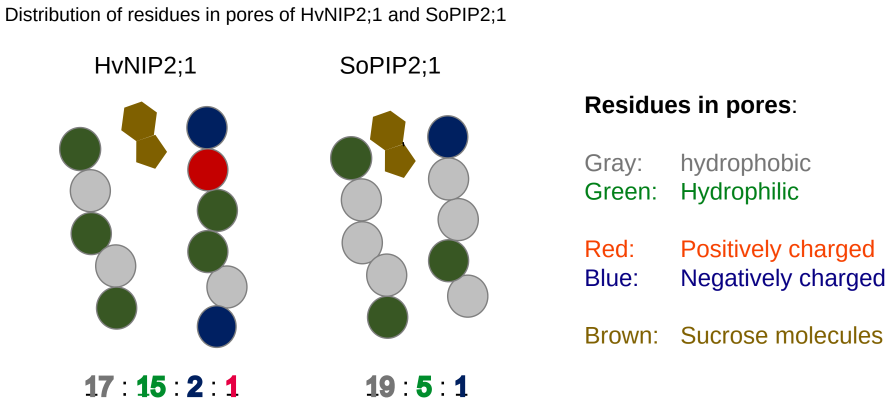


**Fig. S12.** Schematics of residue distributions in the pores of HvNIP2;1 and SoPIP2;1.

Schematics (not drawn to scale) illustrates key features of the pore in HvNIP2;1 that allow sucrose permeation and not that of D-glucose. Hydrophilic environment of the pore with three charged residues (two negatively and one positively charged) and a more voluminous pore in HvNIP2;1 permit sucrose permeation, as explained in Discussion. Shown is also the distribution of residues in the pore of SoPIP2;1, in which sucrose permeation does not occur.
